# Mapping the Genetic Architecture of the Adaptive Integrated Stress Response in *S. cerevisiae*

**DOI:** 10.1101/2024.12.19.629525

**Authors:** Rachel Baum, Jinyoung Kim, Ryan Y. Muller, Nicholas T. Ingolia

## Abstract

The integrated stress response (ISR) is a conserved eukaryotic signaling pathway that responds to diverse stress stimuli to restore proteostasis. The strength and speed of ISR activation must be tuned properly to allow protein synthesis while maintaining proteostasis. Here, we describe how genetic perturbations change the dynamics of the ISR in budding yeast. We treated ISR dynamics, comprising timecourses of ISR activity across different levels of stress, as a holistic phenotype. We profiled changes in ISR dynamics across thousands of genetic perturbations in parallel using CRISPR interference with barcoded expression reporter sequencing (CiBER-seq). We treated cells with sulfometuron methyl, a titratable inhibitor of branched-amino acid synthesis, and measured expression of an ISR reporter. Perturbations to translation such as depletion of aminoacyl-tRNA synthetases or tRNA biogenesis factors reduced cell growth and caused a strikingly proportionate activation of the ISR activation. In contrast, impaired ribosome biogenesis reduced basal ISR activity and weakened ISR dynamics. Reduced ribosome capacity may lower the demand for amino acids and thereby explain these changes. Our work illustrates how CiBER-seq enables high-throughput measurements of complex and dynamic phenotypes that shed light on adaptive and homeostatic mechanisms.

## Introduction

The integrated stress response (ISR) is a conserved, eukaryotic signaling pathway that responds to translational stresses to maintain or restore cellular proteostasis (Costa-Mattioli and Walter 2020). ISR activation leads to the inhibitory phosphorylation of the translation initiation factor eIF2ɑ, which reduces overall protein synthesis, reprograms translation, and indirectly induces adaptive gene expression programs (Costa-Mattioli and Walter 2020). ISR activation must be balanced to enable ongoing protein production while retaining proteostasis; either excessive or inadequate ISR activation can be pathogenic (Delépine et al. 2000; H. P. Harding et al. 2000; Scheuner et al. 2001; Abdulkarim et al. 2015; Kernohan et al. 2015). ISR activation is dynamic, and the signaling pathway can adapt to chronic stress due to feedback at many levels (Novoa et al. 2003; Ma and Hendershot 2003; Klein et al. 2022). Drugs that target these feedback pathways and alter ISR dynamics, such as ISRIB and sephin1, hold particular promise as therapeutics that avoid the toxicity produced by blunt ISR inhibition (Das et al. 2015; Sidrauski et al. 2013; Halliday et al. 2015). Understanding how ISR dynamics change in response to genetic perturbations could shed new light on the ways this pathway maintains productive proteostasis across changing conditions.

The ISR was discovered first as the budding yeast pathway for general amino acid control (GAAC) (Hinnebusch 2005). In response to amino acid deprivation, yeast translationally induce the Gcn4 transcription factor, which in turn activates amino acid biosynthetic genes. The human ISR is activated by a broader range of proteotoxic stressors, but it likewise drives transcriptional induction of genes needed for amino acid synthesis and import (Heather P. Harding et al. 2003). This gene expression program suggests that one major role for the ISR is maintaining adequate levels of amino acids to support protein synthesis. Indeed, uncharged tRNAs are thought to activate the ISR as a direct signal of amino acid insufficiency, although recent evidence suggests an important role for ribosome stalling and collisions as well (Wu et al. 2020). Translation and proteostasis depend on many different aspects of cell physiology, however, potentially connecting them to the ISR.

To understand the budding yeast ISR more broadly, we recently carried out a genome-wide survey to find genetic perturbations that affect ISR activity. We combined CRISPR interference (CRISPRi) knockdown of individual genes with barcoded expression reporters (CiBER-seq) for ISR activity to learn how these knockdowns either activated the ISR in otherwise unstressed cells, or blocked ISR activation during stress (Muller et al. 2020). In addition to identifying known ISR triggers, we found other perturbations of the translational machinery, such as impaired tRNA biogenesis, that activated the ISR. We also found genetic modifiers of ISR activation, such as the known feedback regulator *PCL5*. While *PCL5* depletion changed steady-state ISR activation, we expected that it would show a stronger effect on the dynamics of ISR activation.

We thus set out to find genetic perturbations that changed the strength and timing of ISR activation across different degrees of stress. We used a titratable inhibitor of amino acid biosynthesis, sulfometuron methyl (SM), that provoked adaptive ISR signaling, and profiled the timecourse of ISR activation across different concentrations of SM. We observed genetic perturbations that shifted the sensitivity, onset, and long-term adaptation of the ISR. Most notably, disrupting ribosome biogenesis genes shifted ISR dynamics. Decreased ribosome levels abolished ISR activation at low SM doses and slowed responses at high doses, likely by reducing translational capacity. Our work highlights the importance of amino acid demand in reshaping ISR dynamics.

## Results

### Profiling the adaptive nature of the ISR and selectively screening genetic perturbations that effect these dynamics using CiBER-seq

In budding yeast, the ISR drives adaptive upregulation of amino acid biosynthesis genes in response to amino acid limitation. The herbicide sulfometuron methyl (SM) is a titratable inhibitor of branched amino acid synthesis that activates the ISR in budding yeast (Jia et al. 2000). Lower doses of SM cause transient ISR activation followed by growth recovery, while high doses of SM lead to persistent ISR activation and growth arrest (Jia et al. 2000). Indeed, we found that yeast grown in minimal media showed a dose-dependent growth inhibition upon SM treatment (Fig1A, Supp Fig 1A). To measure ISR activation, we developed a reporter using the native promoter of *PCL5*, a strong transcriptional target of the yeast ISR transcription factor Gcn4 (Shemer et al. 2002). We observed a rapid, dose-dependent induction of endogenous *PCL5* that is mimicked by our citrine reporter driven by the *PCL5* promoter (Fig 1B). High dose (14 µM) SM treatment led to full ISR activation within 15 minutes that persisted over 4 hours. In contrast, low dose (0.56 µM) SM treatment caused slower ISR activation that gradually declined. This dose-dependent, adaptive response offers a setting to learn how genetic changes alter ISR dynamics.

**Figure 1:**
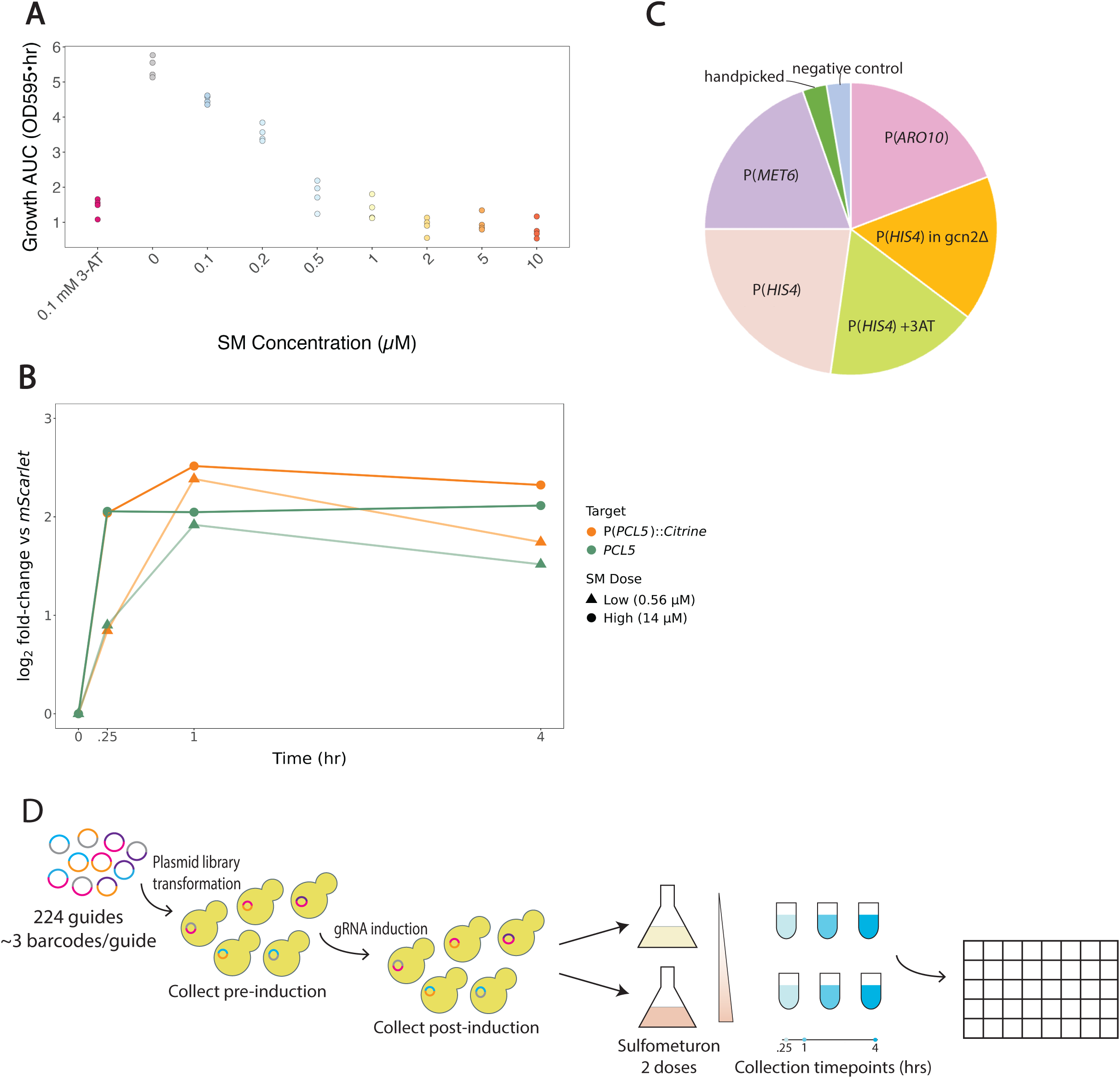
Sulfometuron methyl titratable response and CiBER-seq screen setup. **(A)** Growth AUC (Area Under Logistic Curve) reflects relative growth rate for fitted growth curves for cells maintained in 3-AT or SM. **(B)** RT-qPCR for cells treated with low SM dose (●) or high SM dose (▲) for 0, .25, 1, or 4 hours, normalized to P(*UBC6*)::mScarlet reporter. Endogenous *PCL5* expression shows dose and time-dependent effect. P(*PCL5*)::Citrine reporter expression shows similar dose and time-dependent effect. **(C)** Pie chart of prior CiBER-seq screens guides were chosen from; guides were chosen based on their strength and significance in these screens. **(D)** Schematic of CiBER-seq screen setup. We integrated plasmid libraries containing guides into yeast using a Bxb1 integrase system. We collected the pre-guide induction sample; induced guides with 500 ng/mL tetracycline and collected post-guide induction sample. The culture was divided into two flasks for treatment with low SM dose or high SM dose and cells were collected at 0.25, 1, or 4 hours post-treatment. ISR dynamics were mapped by DESeq2 analysis.

We then set out to profile how genetic perturbations affect ISR dynamics by using the CiBER-seq approach we developed that combines CRISPRi gene knockdown with barcoded expression readout sequencing (Muller et al. 2020). CiBER-seq, and similar techniques, can provide time-resolved measurements of molecular phenotypes, such as ISR reporter expression, across thousands of genetic perturbations in parallel (Muller et al. 2020; Alford et al. 2021). The guide RNAs that induce these genetic perturbations are delivered to cells along with transcriptional reporters that measure their phenotypic effects. The reporter transcript contains a unique, guide-specific barcode sequence, and the RNA abundance of each guide-specific barcode reflects the transcription level of the reporter in cells that contain the associated guide. By sequencing and quantifying expressed RNA barcodes, it is possible to measure reporter expression across many different perturbations in a single, bulk sequencing sample. To correct for variation in the number of cells containing different barcodes, we use two distinct reporters delivered together: the *PCL5* ISR reporter along with a reporter based on the *UBC6* promoter, which is expressed at a very consistent level across many different conditions (Supp Fig 1B). We have found that comparisons between two different expression reporters yields lower background and less technical noise than normalization of expressed reporter RNA against barcode DNA abundance (Muller et al. 2020; Lobel and Ingolia 2024).

We previously used CiBER-seq to profile ISR activity, along with other reporters for amino acid metabolism, in static conditions. Here, we chose over 200 CRISPRi guides that displayed strong and significant effects in prior screens. A majority of guides were selected based on their effect on the classic ISR target *HIS4* either in unperturbed cells or in cells treated with the ISR activating drug 3-AT, with or without deletion of the ISR signaling kinase gene *GCN2* (Muller et al. 2020). Additional guides were chosen because of their effect on *MET6*, which encodes a methionine biosynthesis enzyme (Muller et al. 2020). A final set of guides were chosen due to their effect on a reporter for the catabolic enzyme gene *ARO10* (Diamond, McGlincy, and Ingolia 2024). A small number of guides were added by hand, along with six negative control guides targeting the *HO* endonuclease, for an overall set of 224 guides targeting 185 distinct genes (Fig 1C).

To profile a broader range of perturbations and identify genetic interactions in the ISR, we created a library of pairwise combinations between these guides. Expressing two different CRISPRi guides in the same cell yields strong knockdown of both targets. Importantly, our dual-guide library provided measurements of single-guide phenotypes by pairing each guide with one of several negative controls with no effect. Our library contained 19,265 of the 24,976 possible, distinct guide pairs.

We delivered our dual-guide CiBER-seq library into a yeast strain engineered to facilitate ISR dynamics measurements. Budding yeast is naturally prototrophic for amino acids, but common laboratory strains were rendered auxotrophic to facilitate selective growth. We were concerned that amino acid auxotrophies would distort the ISR, and so we created a strain that would be fully prototrophic in the context of our CiBER-seq library. This strain also expressed the CRISPRi effector (dCas9-Mxi) and the tetracycline repressor (TetR) to provide tetracycline-inducible guide RNA expression (Gilbert et al. 2013; Smith et al. 2016). Finally, it contained a genomic “landing pad” that enables high-efficiency genomic integration mediated by the serine recombinase Bxb1 (Matreyek et al. 2020; Xu and Brown 2016; Lobel and Ingolia 2024). Integration of plasmids from the guide library into this landing pad reconstitutes a split *URA3* marker cassette, allowing selection for integrants and restoring full prototrophy to the strain (Levy et al. 2015; K. Lee, Zhang, and Lee 2008; Lobel and Ingolia 2024). We carried out four independent integrations and collected 3.0x10^5^ – 2.4 x 10^6^ independent transformants.

The transformed populations were grown in minimal media and treated with tetracycline to induce guide expression. We sampled cells before guide expression and then 17.5 hours after guide induction. Each of the cultures were divided in half and treated with either a low dose or high dose of SM. We took samples 0.25 hours, 1 hour, and 4 hours after SM addition. We extracted RNA from each sample, created barcode sequencing libraries from 4 µg RNA per sample, and sequenced the barcode libraries (Fig 1D). We also quantified *PCL5* reporter expression in our libraries by RT-qPCR (Supp Fig 1C). We saw substantial ISR activation even prior to SM treatment, which likely reflects the fact that a large fraction of guides in our library activate the ISR in the absence of external stress.

### Analysis of CiBER-seq screen to identify time-sensitive or dose-sensitive changes in ISR dynamics

We analyzed sequencing data to infer ISR activity and growth phenotypes for each individual genetic perturbation. Guide RNAs that change cell growth will lead to differences in the number of cells harboring that guide RNA, which will in turn change the abundance of the normalizer barcode driven by the housekeeping *UBC6* promoter. Guide RNAs that increase ISR activity will cause higher levels of the *PCL5* reporter barcode, relative to the matched *UBC6* barcode; guides that decrease ISR activity will lower the ratio between these barcodes. We inferred these ISR activity and growth rate jointly at all timepoints in a general linear modeling framework, combining all barcodes for a given genetic perturbation by weighted averaging (Love, Huber, and Anders 2014).

We measured growth and ISR activity phenotypes after guide induction, but prior to SM treatment, that replicated known features of the yeast ISR. Most notably, many genetic perturbations that compromised protein synthesis by reducing the availability of charged tRNA led to slower growth and ISR activation (Fig 2A). Knockdown of aminoacyl-tRNA synthetases (AARS), components of RNA polymerase III (which transcribes tRNAs), and subunits of the tRNA processing enzyme RNase P all showed this effect, along with knockdown of proteins comprising the translation initiation factors eIF2 and eIF2B. There was a striking correlation between ISR activation and growth defect, suggesting that steady-state ISR activity directly tracks the extent to which protein synthesis limits cell growth (Fig 2A). The only exception to this trend was *PCL5*, which encodes a negative feedback regulator of the ISR transcription factor Gcn4 (Shemer et al. 2002). Knockdown of *PCL5* activates the ISR without causing a growth defect, presumably by reducing this feedback loop. Importantly, this *PCL5* guide does not target our ISR reporter, which is based on the *PCL5* promoter. Our library included a second *PCL5* guide that does target the reporter itself as well as the endogenous *PCL5* gene, and this guide strongly reduces reporter expression.

**Figure 2:**
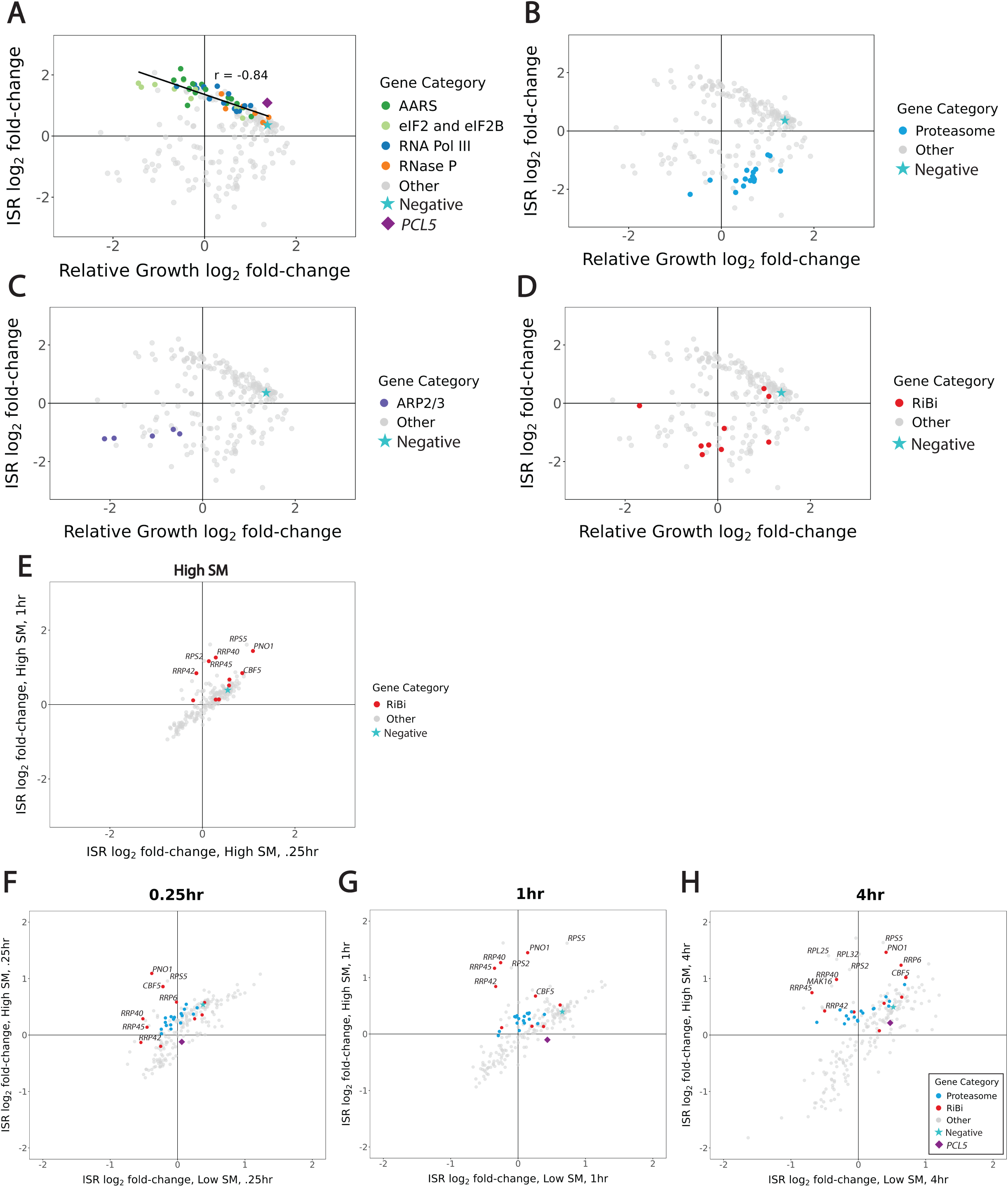
DESeq2 analysis of single mutants show dynamic ISR phenotypes. **(A-D)** Scatterplots show guides targeting functionally related genes have similar phenotypes; ISR log2 fold-change versus growth log2 fold-change after guide-induction **(A)** Guides targeting eIF2 and eIF2B, aminoacyl-tRNA synthetases (AARS), RNA Pol III, and RNaseP components proportionately slow growth and reduces ISR activation. Solid line represents correlation between these guides (r =-0.84). Negative guide (★) is used as a reference point for growth and ISR effects. PCL5 targeted guide (◆) activates the ISR without causing a growth defect. **(B)** Knockdown of proteasome components reduces ISR activity. **(C)** Guides targeting subunits of the ARP2/3 complex reduced cellular fitness and decreased basal ISR signaling. **(D)** Guides targeting Ribosome Biogenesis (RiBi) factors mostly grouped together and led to slower growth and reduced ISR activation. **(E)** Scatterplot of high SM treated cells .25hr versus 1hr shows increase in ISR expression; guides targeting RiBi genes increase ISR activation after 1 hour treatment. **(F-H)** Scatterplots of low versus high dose for each timepoint of SM treatment. Guides targeting ribosome biogenesis increase activation at high SM dose relative to low SM dose at each timepoint. Activation increases over treatment time. Guides targeting proteasome components show a similar but weaker ISR activation.

In contrast, many guides reduced basal ISR activity, with varying effects on growth. Consistent with our earlier CiBER-seq experiments, we saw that knockdown of proteasome components reduced ISR activity (Fig 2B) (Muller et al. 2020). Likewise, guides targeting subunits of the ARP2/3 complex, an essential actin regulator, reduced cellular fitness and decreased basal ISR signaling (Fig 2C). Guides targeting ribosome biogenesis (RiBi) genes led to growth defects and decreased basal ISR signaling (Fig 2D). More broadly, guides targeting functionally related genes tended to show similar phenotypes (Supp Fig 2A). We next analyzed the single-guide ISR phenotypes across timecourses of SM treatment at two different doses. The similarity of dynamic ISR phenotypes between different guides targeting the same gene and between guides with shared function was quite strong.

We were interested in identifying genetic perturbations that change the dynamics of ISR activation. While most genetic perturbations showed a proportionate response to high-dose SM treatment between 0.25 hour and 1 hour, we noticed that several guides targeting ribosome biogenesis factors deviate from this trend and had relatively higher ISR activation at the later timepoint (Fig 2E). Comparison between low and high SM doses at each individual timepoint also demonstrate that most guides show a proportionate response, although more guides lead to dose-dependent differences with increasing length of treatment time. Guides targeting ribosome biogenesis factors again stand out in this comparison, with increased ISR activation at a high SM dose relative to a low SM dose at every treatment timepoint, and increasing over treatment time (Fig 2F-H). Guides targeting the proteasome show a similar, albeit weaker, trend.

We also analyzed dual-guide phenotypes and found that they generally fall within the range delineated by the single guides. In particular, many guide pairs cause proportionate ISR activation and fitness defects (Supp Fig 2B,C). We will present further analysis of these pairwise perturbations elsewhere.

### Ribosome biogenesis transcripts *RRP42* and *PNO1* regulate the ISR in a dose-dependent manner

While we observed a wide range of ISR activity phenotypes, knockdown of ribosome biogenesis factors stood out for changing ISR dynamics in a dose- and time-dependent manner. We wished to understand the basis for this distinctive phenotype. Perturbations of ribosome biogenesis genes *RRP40*, *RRP42*, *RRP45*, *PNO1* and *CBF5* led to similar fitness defects and reduction in basal ISR activity (Fig 3A). We validated the effect of *RRP42* knockdown levels on cell fitness by titrating the concentration of tetracycline, which regulates the knockdown level. Knockdown efficiency had an inverse relationship with the growth rate, highlighting the importance of the ribosome biogenesis factor *RRP42* in maintaining cell fitness (Supp Fig 3A,B). Across all five ribosome biogenesis genes, ISR activity remained low when yeast were exposed to low doses of SM, but high doses led to strong but delayed ISR activation (Fig 3B). *RRP40*, *RRP42*, and *RRP45* encode components of the RNA exosome, a multifunctional RNA exonuclease that processes pre-rRNAs (Kilchert, Wittmann, and Vasiljeva 2016; Okuda et al. 2020). *PNO1* and *CBF5* both encode dedicated ribosome biogenesis factors involved in ribosomal RNA processing (Senapin et al. 2003; Lafontaine et al. 1998). We carried out targeted validation experiments to confirm the phenotypes we measured for these guides in our screen. Knockdown of *RRP42* following SM treatment confirmed that perturbed cells did not greatly activate the ISR after a low-dose SM treatment, but did reach nearly wildtype levels after 1 hour of high-dose treatment (Fig 3C and Supp Fig 3C,E,G). Knockdown of *PNO1* likewise impaired ISR activation at a low SM dose that is overcome when cells are treated with a high SM dose—surpassing wildtype ISR activation (Fig 3D, Supp Fig 3D,F,H). Knockdown of *CBF5* showed a similar pattern of weak ISR activity at low SM doses with robust responses at a higher dose (Supp Fig 4A-D).

**Figure 3:**
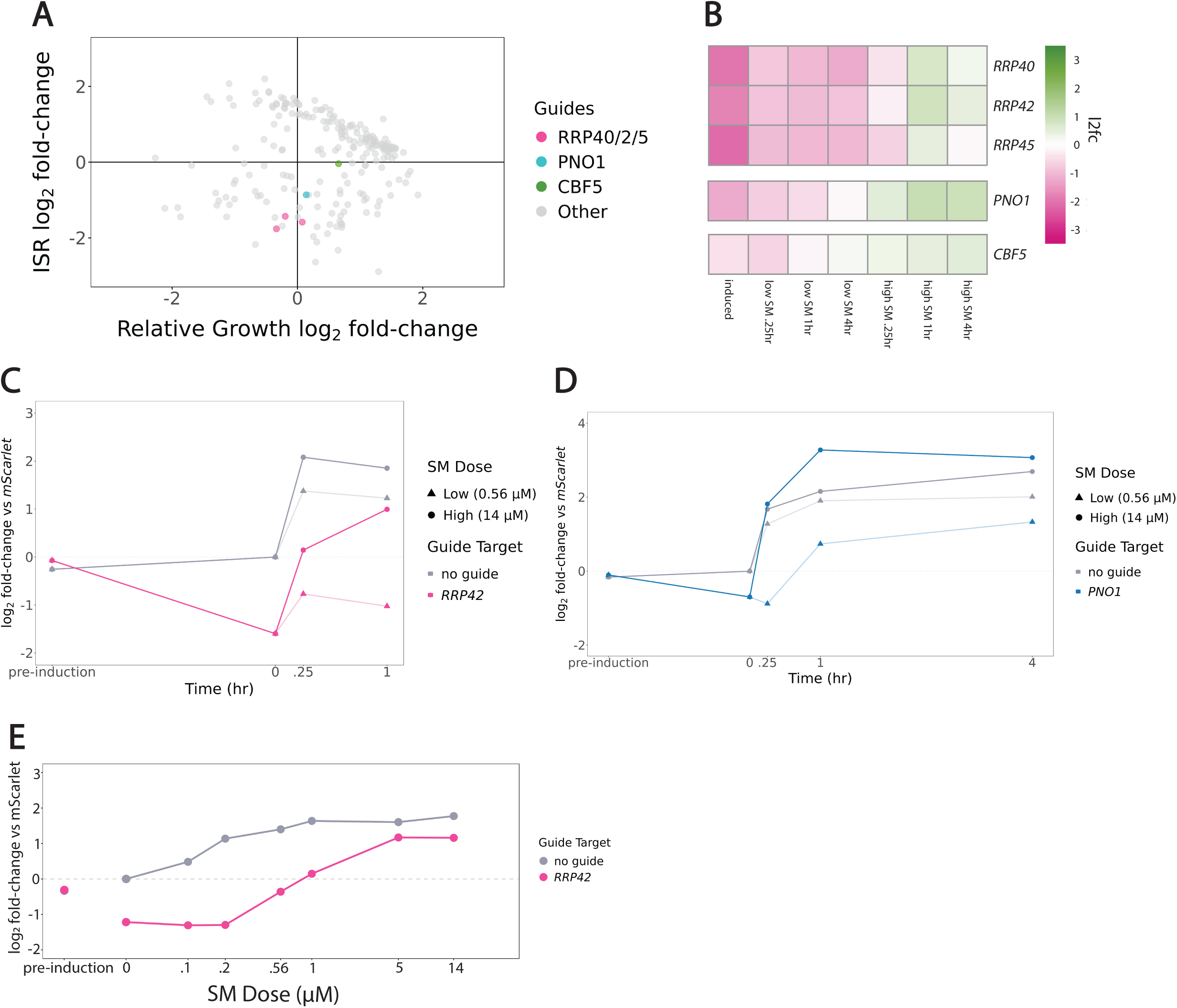
Knockdown of RiBi factors *RRP42* and *PNO1* lead to SM treatment dose-dependent response. **(A)** Scatterplot highlighting *RRP40/2/5, PNO1*, and *CBF5* targeted guides. Guides targeting these RiBi genes slows cell growth and decreases ISR activation. **(B)** Heatmap for *RRP40*, *RRP42*, *RRP45*, *PNO1*, and *CBF5* mutants shows ISR log2 fold-change normalized to negative guide reference. Reduced ISR activity after low SM dose treatment; high ISR activity after high SM dose treatment. **(C)** RT-qPCR of ISR reporter for wildtype and *RRP42* knockdown cells, replicate 1. ISR reporter normalized to mScarlet for pre-guide induction, post-guide induction and 0, .25, and 1 hour low or high SM treated cells. Low SM dose reduces ISR activation and high SM dose strengthens ISR activation. **(D)** RT-qPCR of ISR reporter for wildtype and *PNO1* knockdown cells, replicate 1. ISR reporter normalized to mScarlet for pre-guide induction, post-guide induction and 0, .25, 1, and 4 hours low or high SM treated cells. Low SM dose reduces ISR activation and high SM dose strengthens ISR activation. **(E)** RT-qPCR for ISR reporter in wildtype or *RRP42* knock-down cells in pre-guide induction, post-guide induction, and SM treatment doses after 1 hour of treatment. *RRP42* knockdown decreases ISR SM response.

**Figure 4:**
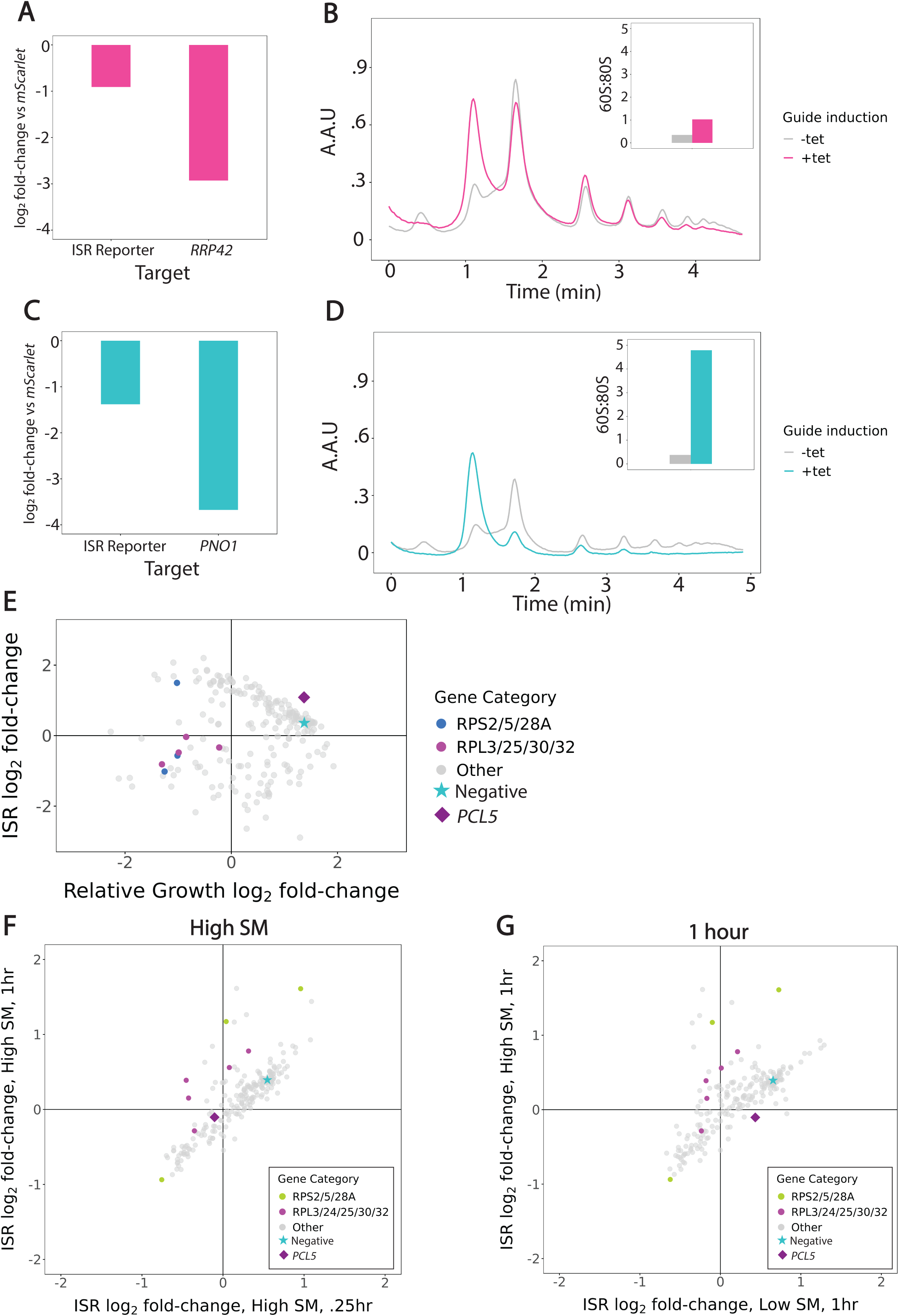
Knockdown of *RRP42* and *PNO1* leads to defects in 40S ribosome biogenesis that reshape the dynamics of the ISR. **(A)** RT-qPCR data pre- and post-guide induction for RRP42 knockdown lysate. *RRP42* knock-down reduced ISR reporter expression; *RRP42* shows robust knock-down efficiency. **(B)** 10-50% sucrose gradients in pre and post guide tet-induction in *RRP42* cells. Time (min) versus Absorbance Arbitrary Units (A.A.U). 60S:80S ratio for -/+ knockdown shows increase in ratio after knockdown. **(C)** RT-qPCR data pre- and post-guide induction for PNO1 knockdown lysate. PNO1 knock-down reduced ISR reporter expression; *PNO1* shows robust knock-down efficiency. **(D)** 10-50% sucrose gradients in pre- and post guide tet-induction in *PNO1* cells. Time (min) versus Absorbance Arbitrary Units (A.A.U). 60S:80S ratio for -/+ knockdown shows increase in ratio after knockdown. **(E)** Scatterplot highlighting guides targeting small and large ribosomal subunits shows strong growth defect and mild reduction of ISR activation. **(F)** Scatterplot highlighting guides targeting small and large ribosomal subunits after high SM dose treatment at .25hr versus 1hr shows similar dose-dependent ISR effect. **(G)** Scatterplot highlighting guides targeting small and large ribosomal subunits after 1hr low versus high dose SM treatment shows similar time-dependent ISR effect.

To further characterize how *RRP42* knockdown affects the response to SM, we measured ISR activation at 1 hour across a range of concentrations. *RRP42* knockdown cells show decreased basal ISR activity that does not increase below a threshold of 0.2 µM SM, whereas wildtype cells induce the ISR at these doses. Overall, *RRP42* knockdown cells appear 4– 5-fold less sensitive to SM (Fig 3E). We observed a similar trend when measuring the activation of the endogenous ISR target *HIS4* (Supp Fig 3I).

### Cellular defects in 40S ribosome biogenesis reshape the dynamics of the ISR

To directly measure the impact of *RRP42* and *PNO1* knockdown on translation, we analyzed polysomes in these cells. We generated matched lysates from knockdown and control cells (Fig 4A,C) and performed polysome profiling by sucrose density gradient ultracentrifugation. Both *RRP42* and *PNO1* knockdown demonstrated a loss of 40S subunits in comparison with wildtype cells (Fig 4B,D). *RRP42* knockdown led to some loss of polysomes and an increase in free 60S subunits (Fig 4B). *PNO1* knockdown caused stronger polysome collapse and a dramatic increase in the 60S:80S suggesting reduced overall translation (Fig 4D). We validated the loss of 40S subunits on shallower density gradients with better resolution of individual subunits (Supp Fig 5A-B).

**Figure 5:**
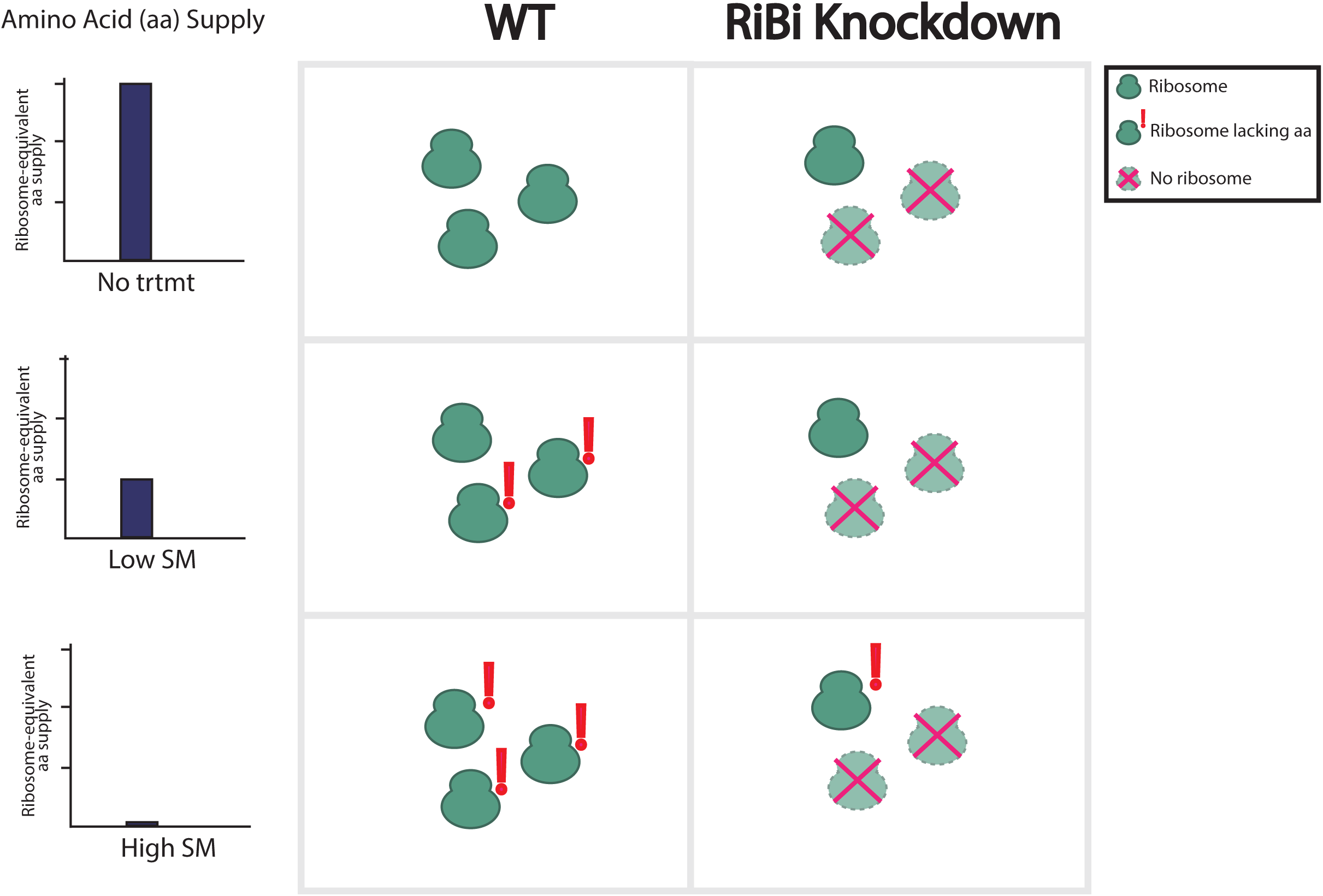
Model of the effect of ribosome levels on reshaping ISR dynamics. Ribosome biogenesis mutants decrease the availability of ribosomes, reducing cellular demand for amino acids. **Low SM dose**: Wildtype cells can maintain amino acid levels by activating the ISR and inducing amino acid biosynthetic genes. Ribosome biogenesis mutants have weaker demand for amino acids due to lower ribosome availability, leading to weaker and slower ISR activation. **High SM dose**: Ribosome biogenesis mutants do become limited for amino acids, which leads to higher ISR activation in a manner similar to wildtype cells.

Since these polysome traces demonstrate that defects in 40S ribosome biogenesis reshape the dynamics of the ISR, we turned to our dataset to understand whether this effect was specific to the small ribosomal subunit. We evaluated guides targeting protein components of the 40S subunit (*RPS2*, *RPS5*, and *RPS28A*) and the 60S subunit (*RPL3*, *RPL25*, *RPL30*, and *RPL32*). In nearly all cases, knockdown of these subunits showed a strong growth defect and mild reduction in ISR activation (Fig 4E). Ribosomal protein depletion did not show the same strong ISR dynamics phenotype that we observed from disrupting ribosomal RNA processing. While we do observe that ribosomal protein alters dose-dependent ISR activation at 4 hours, this phenotype is not clearly different between guides targeting the small and large subunits (Fig 4F-G). This suggests that this is not a subunit specific response, but more likely a consequence of limited ribosome availability.

## Discussion

We identify genetic perturbations that distort the behavior of the ISR. We treated ISR dynamics—the timecourse of ISR activation across different degrees of stress—as a holistic phenotype and profiled ISR dynamics across thousands of genetic perturbations in parallel using CiBER-seq. By perturbing genes known to affect the ISR, we learned how each of these individual genes help shape the dynamics of this response.

Our results point to a relationship between ribosome levels, amino acid demand and ISR activation. We propose that ribosome levels impact ISR dynamics—ribosome biogenesis mutants decrease the availability of ribosomes, reducing cellular demand for amino acids. At a low SM dose, wildtype cells can maintain amino acid levels by activating the ISR and thereby inducing amino acid biosynthetic genes. In contrast, ribosome biogenesis mutants have weaker demand for amino acids due to lower ribosome availability, leading to weaker and slower ISR activation. At a high SM dose, ribosome biogenesis mutants do become limited for amino acids, which leads to higher ISR activation, ultimately reaching levels similar to those seen in wildtype cells (Fig 5).

Disrupting ribosome biogenesis has many effects on the cell. On short timescales, ribosome biogenesis defects lead to aggregation of unassembled ribosomal proteins and proteotoxic stress (Tye et al. 2019; Albert et al. 2019). On longer timescales, ribosome deficiency leads to widespread secondary effects on gene expression (Cheng et al. 2019). Changes in ISR sensitivity reflect another feature of this altered cellular environment.

ISR dynamics comprise a multi-dimensional phenotype. We inferred ISR activity across 7 distinct measurements, providing several degrees of freedom for different genetic perturbations to change these measurements. In addition to the slow and weak ISR associated with ribosome biogenesis defects, we also observed changes to adaptation after prolonged exposure. For example, perturbing *SWM2*, which has a role in snRNA/snoRNA cap trimethylation, or *MOT1*, involved in transcriptional regulation, only shifted ISR dynamics at the late 4 hour timepoint of treatment, indicating a possible effect on adaptation. Understanding these long-term effects could shed light on the trade-offs and limits in long-term ISR adaptation.

## Acknowledgements

We thank Joseph Lobel and all members of the Ingolia lab for thoughtful discussion and commentary. This work was supported by the National Institute of General Medical Sciences of the National Institutes of Health under award numbers R01 GM135233 (N.T.I.) and by the National Science Foundation Graduate Research Fellowship under Grant No. DGE 1752814 (J.K.). Any opinions, findings, and conclusions or recommendations expressed in this material are those of the authors and do not necessarily reflect the views of the National Science Foundation. Short-read sequencing was performed at the UCSF CAT, supported by UCSF PBBR, RRP IMIA, and NIH 1S10OD028511-01 grants. Long read sequencing was carried out at the DNA Technologies and Expression Analysis Cores at the UC Davis Genome Center, supported by NIH Shared Instrumentation Grant 1S10OD010786-01.

## Methods

### Yeast Strains

Strains used in this study are listed in Table 1. Strains were derived from *S. cerevisiae* BY4741 using standard genetic techniques and CRISPR-Cas9 technology (M. E. Lee et al. 2015). Yeast transformations were carried out using standard LiAc/PEG protocol and grown at 30 °C (Gietz and Schiestl 2007). Strain with dCas9-Mxi, TetR, and attP genomic landing pad was previously described in (Lobel and Ingolia 2024; Muller et al. 2020). Yeast were rendered prototrophic by re-integration of wildtype *HIS3* and *MET15* (Mülleder et al. 2012); integrated landing pad plasmid pNTI829 contained the *K. lactis LEU2* gene, and integration of donor plasmids into the landing pad reconstituted *S. cerevisiae URA3* with a synthetic intron.

**Table 1:** Yeast strains used in this study. All strains are derived from *S. cerevisiae* BY4741.

| Strain Number | Genotype | Source |
| --- | --- | --- |
| NIY539 | S288C MATa HIS3 leu2Δ0 LYS2 MET15 ura3Δ0 XII-2:kanMXsyn:dCas9-Mxi1:TetR X-3:KILEU2:P(URA3)::URA3N-attP-2A-Bxb1-V5::T(ADH1) | Lobel et al. 2024 |
| NIY559 | NIY539 X-3:KILEU2:P(URA3)::URA3N-attP-2A-Bxb1-V5::T(ADH1)::Ura3N/C P(Pcl5)-Citrine/p(UBC6)-mScarlet | This work |
| NIY575 | NIY539 X-3:KILEU2:P(URA3)::URA3N-attP-2A-Bxb1-V5::T(ADH1)::Ura3N/C P(Pcl5)-Citrine/p(UBC6)-mScarlet X-4::hphMX:sg(RRP42) | This work |
| NIY585 | NIY539 X-3:KILEU2:P(URA3)::URA3N-attP-2A-Bxb1-V5::T(ADH1)::Ura3N/C P(Pcl5)-Citrine/p(UBC6)-mScarlet [sg(PNO1)] | This work |
| NIY587 | NIY539 X-3:KILEU2:P(URA3)::URA3N-attP-2A-Bxb1-V5::T(ADH1)::Ura3N/C P(Pcl5)-Citrine/p(UBC6)-mScarlet X-4::hphMX:sg(CBF5) | This work |

Strains carrying individual gRNAs were integrated using either the Bxb1 recombinase system or homologous recombination into the X-4 locus using the Easyclone 2.0 vector genomic integration system (Matreyek et al. 2020; Matreyek, Stephany, and Fowler 2017; Durrant et al. 2023; Lobel and Ingolia 2024; Stovicek et al. 2015). All strains used for CiBER-seq screen and gRNA validation experiments were prototrophic and grown in minimal media. Minimal media composition: Yeast nitrogen base without amino acids (Thomas Scientific #C994M99) + 2% dextrose.

### Plasmids

Plasmids used in this study are listed in Table 2. Divergent promoter reporter plasmids with guides were made using Gibson-style NEB HiFi assembly (NEB #E2621L) (Gibson et al. 2009). Individual guide plasmids in Easyclone vectors were made using Gibson-style cloning into Easyclone 2.0 vectors with NEB HiFi assembly (Gibson et al. 2009). Plasmids were validated with whole plasmid sequencing performed by Plasmidsaurus using Oxford Nanopore Technology with custom analysis and annotation.

**Table 2:**
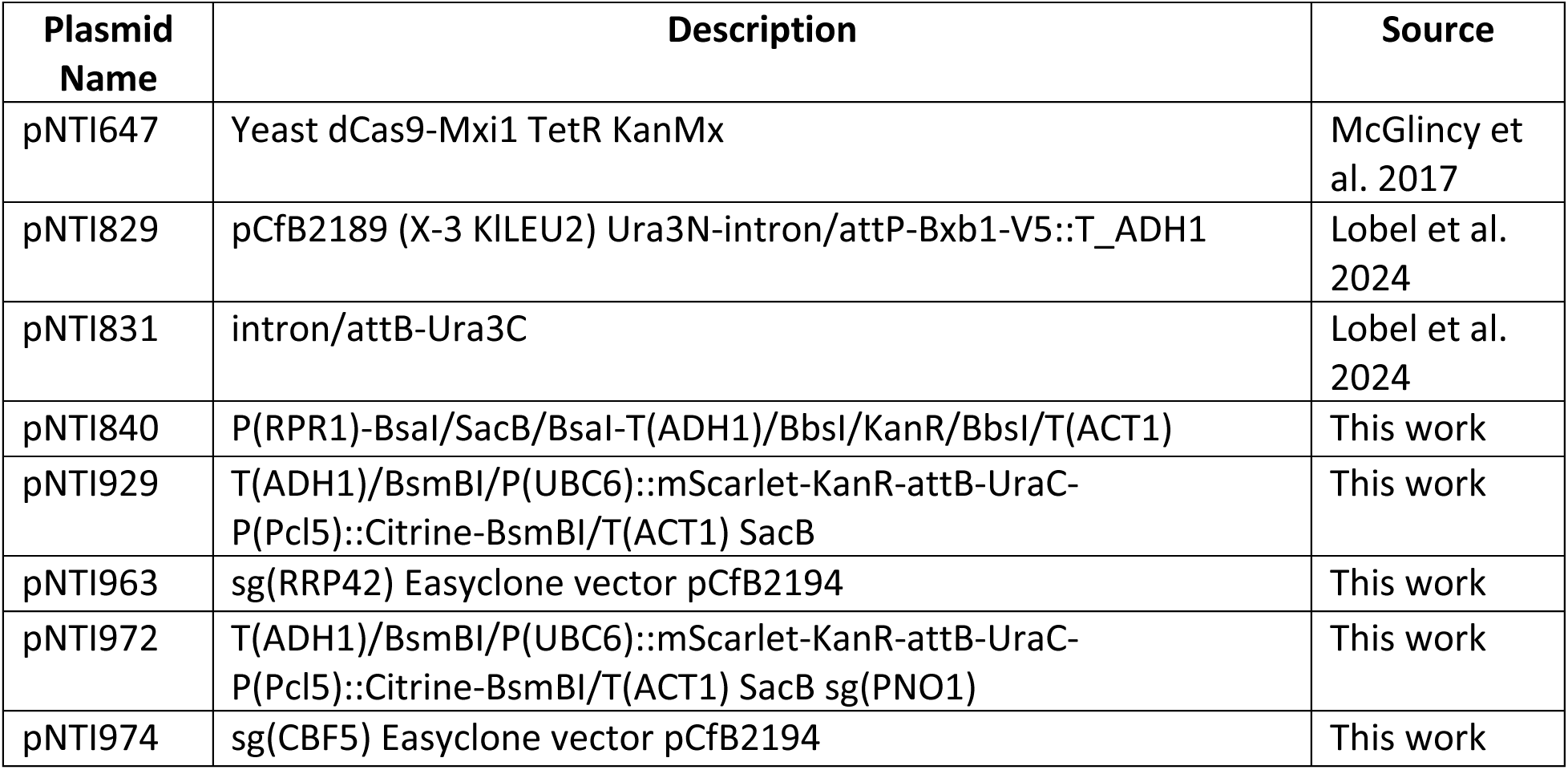
Plasmids used in this study.

| Plasmid Name | Description | Source |
| --- | --- | --- |
| pNTI647 | Yeast dCas9-Mxi1 TetR KanMx | McGlinicy et al. 2017 |
| pNTI829 | pCfB2189 (X-3 KILEU2) Ura3N-intron/attP-Bxb1-V5::T_ADH1 | Lobel et al. 2024 |
| pNTI831 | intron/attB-Ura3C | Lobel et al. 2024 |
| pNTI840 | P(RPR1)-BsaI/SacB/BsaI-T(ADH1)/BbsI/KanR/BbsI/T(ACT1) | This work |
| pNTI929 | T(ADH1)/BsmBI/P(UBC6)::mScarlet-KanR-attB-UraC-P(Pcl5)::Citrine-BsmBI/T(ACT1) SacB | This work |
| pNTI963 | sg(RRP42) Easyclone vector pCfB2194 | This work |
| pNTI972 | T(ADH1)/BsmBI/P(UBC6)::mScarlet-KanR-attB-UraC-P(Pcl5)::Citrine-BsmBI/T(ACT1) SacB sg(PNO1) | This work |
| pNTI974 | sg(CBF5) Easyclone vector pCfB2194 | This work |

### Library guide choice

Script for picking guides for this CiBER-seq screen is available upon request. Most guides were chosen from previous screens of *HIS4*, *MET6* or *ARO10* reporters based on the strength of response and significance value. A small number of guides were handpicked. Six negative control guides targeting the *HO* endonuclease. In total there are 224 guides targeting 185 distinct genes. Guide sequences are listed in Table 4.

We excluded some guides from downstream interpretation based on our analysis of the GC content of the protospacer sequence. In general, low GC content in guides leads to more off-target effects, with spacer sequences requiring an optimal GC content of 40-60% (Konstantakos et al. 2022). After identifying one guide with phenotypic effects that seemed unrelated to the designed target (YIR013C_01), we identified all guides with <25% GC (Supp Figure 1D). GC content for each guide is listed in Table 4.

### Library construction

To construct the dual-guide library, we first constructed two sub-pools of single-guide libraries, “pool A” with 120 guides and “pool B” with 100 guides. Oligonucleotide pools (IDT) containing 60 nt protospacers were amplified in 2 x 50. µl reactions with 25. µl 2x Q5 master mix, 2.5 µl oligos NI-1181 and NI-1182 at 10 µM each, 0.5 µl oligo pool at 10 nM, and 22. µl water. PCR conditions were 30 s at 98 °C followed by 12 cycles of 5 s at 90 °C, 20 s at 55 °C, and 10 s at 72 °C, followed by 2 min at 72 °C. products were purified using a Zymo Clean and Concentrator-5, using 500 µl binding buffer with 100 µl pooled reaction and eluted into 6.0 µl elution buffer, yielding 100–120 ng / µl. Products were confirmed by High Sensitivity D1000 assay (Agilent # 5067-5584 and 5067-5585) on an Agilent TapeStation. Vector plasmid was prepared by digesting 500 ng of plasmid DNA in a 50. µl reaction with 1x CutSmart buffer and 1.0 µl BsaI-HFv2 (NEB # R3733S), incubated at 37 °C for 20 hours. Digested vector was purified using a Monarch PCR & DNA Cleanup Kit (5 µg) (NEB # T1030) with 100 µl binding buffer and elution into 6.0 µl. Assembly reactions comprised 100 ng (∼35 fmol) vector and ∼70 fmol amplified guides in a 5.0 µl volume with 5.0 µl 2x HiFi assembly mix, incubated for 1 hour at 50 °C. Assemblies were purified using a Monarch PCR & DNA Cleanup Kit (5 µg) by adding 40. µl water to each assembly followed by 100 µl binding buffer, with a 6.0 µl elution volume. Each assembly was transformed into chemically competent XL1-Blue *E. coli* and allowed to recover 1 hour at 37 °C in 900 µl SOC media. A small aliquot was taken to plate serial dilutions on LB chloramphenicol plates and grown overnight at 37 °C, while the bulk of the transformation was used to 25 ml LB chloramphenicol (35 µg / ml) and grown at 30 °C for 20 hours with shaking. Selective cultures were transferred to 37 °C and grown to a final OD of ∼2.0, and 15 OD•ml units of each culture were harvested by centrifugation at 3,000 x g for 5 minutes. Plasmid DNA was isolated using a Monarch Plasmid Miniprep Kit (NEB #T1010) with 30. µl elution volume, yielding 0.5–2.1 µg DNA per library.

Dual-guide libraries were constructed by digesting 500 ng of each single-guide library in a 50. µl reaction with 1x NEBuffer r3.1 and 1.0 µl BsmBI-v2 (NEB # R0739S) at 55 °C for 16 hours. A 1.5 kb PCR amplicon containing the KanR marker was amplified in a 50 µl reaction with 25. µl 2x Q5 master mix, ∼1.0 pg pNTI840, and 25 pmol each NI-1253 and NI-1254. PCR conditions were 30 s at 98 °C, followed by 30 cycles of 5 s at 98 °C, 10 s at 63 °C, and 45 s at 72 °C, followed by 2 min at 72 °C. Digestions and PCR were purified using a Monarch PCR & DNA Cleanup Kit (5 µg) with 250 µl binding buffer and 6.0 µl elution volume. Four dual-guide sub-libraries were constructed by assembling each pairwise combination of “first” and “second” single-guide sub-libraries. Assemblies used 35 fmol each of “first” and “second” single-guide sub-library digest and 70 fmol of KanR PCR amplicon in a 5.0 µl volume with 5.0 µl 2x HiFi assembly mix, incubated 1 hour at 50 °C. Assemblies were purified using a Monarch PCR & DNA Cleanup Kit (5 µg) by adding 40. µl water to each assembly followed by 100 µl binding buffer, with a 6.0 µl elution volume. Assemblies (1.5 µl of each) were transformed into NEB 10-beta Electrocompetent E. coli (NEB # C3020K) following manufacturer instructions, with 1 hour outgrowth at 37 °C. A small aliquot was taken to plate serial dilutions on LB Kan plates and grown overnight at 37 °C, while the bulk of the transformation was inoculated into pre-warmed LB Kan and grown overnight at 30 °C. Cultures reached an OD of ∼4.0, and 10 OD•ml units were harvested by centrifugation at 16,000 x g for 30 seconds and plasmid DNA was isolated using a Monarch Plasmid Miniprep Kit with 30. µl elution volume, yielding ∼600 ng DNA per library.

Barcodes were introduced by digesting 400 ng DNA from each dual-guide sub-library in a 50. µl reaction with 1.0 µl CutSmart buffer and 1.0 µl BbsI-HF (NEB # R3539S) for 20 hours at 37 °C. Digestions were purified using a Monarch PCR & DNA Cleanup Kit (5 µg) with 100 µl binding buffer and 6.0 µl elution volume. A PCR amplicon containing the AmpR marker was amplified in a 50 µl reaction with 25. µl 2x Q5 Hot Start master mix (NEB #M0494L), 25 pmol each NI-1170 and NI-1171, and 1 pg pNTI831 template. PCR conditions were 30 s at 98 °C, followed by 30 cycles of 5 s at 98 °C, 10 s at 61 °C, and 35 s at 72 °C, and a final 2 min extension at 72 °C. The PCR was purified using a Monarch PCR & DNA Cleanup Kit (5 µg) with 250 µl binding buffer and 20.0 µl elution volume. The purified amplicon was then digested in a 50 µl reaction with 1x CutSmart and 1.0 µl DpnI (NEB #R0176S) for 20 hours at 37 °C to eliminate template and purified again using a Monarch PCR & DNA Cleanup Kit (5 µg) with 250 µl binding buffer and 10.0 µl elution volume. Barcodes were added in an assembly reaction with 140 fmol digested sub-library, 140 fmol PCR amplicon, and 10 pmol each NI-1169 and NI-1172 in a 10.0 µl volume, combined with 10.0 µl 2x HiFi assembly mix and incubated 1 hour at 50 °C. Assemblies were purified using a Monarch PCR & DNA Cleanup Kit (5 µg) by adding 30. µl water to each assembly followed by 100 µl binding buffer, with a 6.0 µl elution volume. Purified assemblies were transformed by combining 4.0 µl DNA with chemically competent XL1-Blue cells, incubating 20 minutes on ice, transforming by heat shock for 30 s at 42 °C, recovering cells on ice with 900 µl SOC for 2 minutes, followed by 1 hour of outgrowth at 37 °C. A small aliquot of cells were taken and used to plate serial dilutions on LB Carb plates grown overnight at 37 °C. The bulk of the transformation was used to inoculate 20 ml of LB Carb liquid media and grown overnight at 30 °C, then transferred to 37 °C and grown to reach an OD of ∼1.0. Cells from two 12 OD•ml of samples of each culture were collected by centrifugation at 3,000 x g for 10 minutes and plasmid DNA was isolated using a Monarch Plasmid Miniprep Kit with 30. µl elution volume. Both samples of each individual sub-library were pooled with a total yield of 9– 22 µg of DNA per sub-library. Estimated barcode diversity for each sublibrary was: P had 87,000 barcodes for 14,000 guide pairs; Q had 30,000 barcodes for 12,000 guide pairs; R had 42,000 barcodes for 12,000 guide pairs; and S had 49,000 barcodes for 10,000 guide pairs.

We cloned the divergent P(*UBC6*)-mScarlet and P(*PCL5*)-citrine reporters into the barcoded libraries in four separate pools. For the barcoded-libraries, we digested 2 µg with BsmBI-v2 (NEB #R0580) overnight at 44 °C. For reporter vector (pNTI929), we digested 2.7 µg with BsmBI-v2 (NEB #R0580) overnight at 44 °C. The digested barcoded-libraries and reporter constructs were purified with the Zymo clean and concentrator kit (Zymo #D4004) according to the manufacturers protocol. We performed a HiFi assembly (NEB #E2621L) with 150 fmole of the digested barcoded-libraries and 304 ng of the digested reporters at 50℃ for 1 hour. This reaction was purified with the Zymo clean and concentrator kit (Zymo #D4004) and eluted in 6µl of water. Each of these assemblies was electroporated by adding 2 µl of the Gibson assembly into 2 x 25 µl MegaX DH10B T1R Electrocomp cells (Thermo Fisher #C640003) according to the manufacturer’s protocol. Cells were recovered by adding 975 µl of SOC media and shaken at 37℃ for 1 hour. Serial dilutions of transformations were plated to ensure sufficient library diversity (>100 unique transformants for each barcode). Culture was transferred into 400 mL LB-Kan and grown overnight to final OD_600_ of 1.8-2.5. Transformation efficiencies ranged from 3.1x10^7^ – 1.0x10^8^. Each of the CiBER-seq libraries were purified using Qiagen HiSpeed midiprep kit (Qiagen #12143).

### PacBio

We prepared a equimolar mix of barcodes from the four dual-guide sublibraries, comprising 4 µl DNA in total, and split this into 2 x 50. µl reactions with 1x rCutSmart, one digested with 1.0 µl EagI-HF (NEB # R3505S) and the other with 1.0 µl SbfI-HF (NEB #R3642S). Digests were incubated for 2.5 hours at 37 °C and DNA was purified using a Monarch PCR & DNA Cleanup kit with 100 µl binding buffer and 10.0 µl elution volume. Digests were pooled and submitted for high-accuracy, long-read circular consensus sequencing. The resulting data were analyzed to extract barcode and guide protospacer sequences using our custom tools designed for this purpose (Kim et al. 2024).

### CiBER-seq screen

Each plasmid sub-library was transformed individually into NIY539 using standard lithium acetate transformation (Gietz and Schiestl 2007). Parental NIY539 containing the attP landing pad was grown overnight to saturation in YEPD. Yeast were diluted to OD_600_ 0.1 into 1l YEPD. Cells were grown up for 5.5 hours to mid-log phase (OD_600_ 0.5–0.7). Cells were collected by centrifugation at 3,000 x g for 5 minutes and washed 3x with water. Following the standard LiAc/PEG transformation protocol, 20 µg of each CiBER-seq library was transformed with heat shock at 42 °C for 40 minutes, mixing by inversion every 5 minutes. Cells were collected by centrifugation at 3,000 x g for 5 minutes and resuspend in 2 mL SCD -Ura media. A small sample was taken to plate serial dilutions on SCD -Ura to assess transformation efficiency and ensure sufficient library diversity (≥10X coverage of library unique barcodes). The bulk of the transformation was transferred to 450 mL SCD -Ura media and grown at 22 °C until it reached an OD_600_ of ∼1.0. After reaching target density, each sub-library population was individually harvested by centrifugation and stored at -80 °C in YEPD + 15% glycerol. The liquid cultures corresponded to 3.0x10^5^ – 2.4 x 10^6^ total transformants.

Yeast populations transformed with plasmid libraries were inoculated at a ratio determined by the number of distinct barcodes in each sub-library (9P:3Q:4R:5S) for a total of 200 OD units. Cells were combined, collected by centrifugation at 3,000 x g for 5 minutes, washed in 10 mL minimal media, and then resuspended in 5 mL minimal media. Cells were transferred into 200 mL prewarmed minimal media and grown for 5 hours at 30 °C until OD_600_ of ∼1.5–2. Cells were then back-diluted to OD_600_ of 0.05 for 5.5 hours until OD ∼0.10–0.15. Pre-induction samples from 50 ml culture were harvested by centrifugation at 3,000 x g for 5 min and stored at -80 °C. Cultures were then diluted to OD_600_ of 0.05 in minimal media with 500 ng / ml tetracycline to induce guides. After cells reached an OD_600_ of 0.7, post-induction samples were harvested. Cells were then divided into two flasks and treated with 0.56 µM (low dose) or 14. µM of sulfometuron methyl (Millipore Sigma #34224). Samples were taken at .25 hours, 1 hour, and 4 hours after sulfometuron methyl addition.

RNA was isolated from each cell sample using an amended protocol from (Ares 2012). Cell pellets were resuspended in 400 µl AE buffer (50 mM NaOAc, pH 5.2, 10 mM EDTA), 40 µl 10% SDS, and 400µl Phenol:Chloroform:Isoamyl alcohol (Fisher Scientific #bp1752i-100) and then incubated at 65 °C with mixing at 2,000 rpm for 15 minutes (Eppendorf ThermoMixer-C). Samples were incubated on ice for 5 minutes and phases were separated by centrifugation at 14,000 rpm at 4 °C for 5 min. We did two washes with 400 µl of chloroform, by adding a layer, shaking the tube and centrifuging at 14,000 rpm at 4 °C for 5min. The aqueous phase was transferred to a new tube, and 50µl of 3M NaOAc and 1µl glycoblue (Invitrogen #AM9516) were added, followed by 100% ethanol until tube was filled. Tubes were mixed by inversion, centrifuged at 14,000 rpm at 4 °C for 20 min. The supernatant was removed and the RNA pellet was washed with 70% ethanol and recollected by centrifugation at 14,000 rpm at 4 °C for 5min and the supernatant was removed. The pellets were air dried for 5 minutes and resuspended in water.

We treated 20 µg of RNA with Turbo DNase (Thermo Fisher Scientific #AM2239) for 30 minutes at 37 °C. RNA was isolated using an RNA Clean and Concentrator-25 kit (Zymo #R1017) according to the manufacturers protocol and eluted in 30 µl water. Reverse transcription was carried out using 4 µg of RNA template with the ProtoScript II reverse transcriptase (NEB #M0368L) and 0.25 µM gene-specific priming (P(*UBC6*)-mScarlet: NI-1267; P(*PCL5*)-citrine: NI-1268) according to manufacturer’s protocol with reaction volume scaled 20-fold. Reverse transcription was carried out at 42 °C for 1hour and inactivated at 80 °C for 5 min. RT reactions were purified using a Zymo Clean and Concentrator-5 kit (Zymo #D4004) with 7:1 ratio of binding buffer to sample, as recommended for ssDNA. Round I PCR was performed using Q5 High-Fidelity 2X Master Mix with primers NI-1272/NI-1273 for the citrine reporter and NI-1271/NI-1273 for the mScarlet reporter. Primers are listed in Table 3. The PCR was scaled up to 20 reactions and run 7 cycles for citrine amplicons and 10 cycles for mScarlet amplicons with Tm 68 °C. The PCR reaction was purified using the Zymo Clean and Concentrator-5 kit with 5:1 protocol and eluted in 15µl water. We quantified the first round PCR product by qPCR using primers NI-827/NI-828 and a diluted series of a standard. We did the second round of PCR using 1000 pM of PCR-I product for 7 cycles with dual indexes (provided by UCSF CAT). Libraries purified using the Zymo Clean and Concentrator-5 kit with 5:1 protocol. Samples were quantified using Qubit 1X dsDNA HS Assay Kit (Thermo Scientific #Q33230) and Agilent Tapestation 2200 instrument. Libraries were pooled and concentrated using PCR Cleanup Beads (UCB DNA Sequencing Facility). Pooled libraries were sequenced on an Illumina Novaseq-X with paired-end 150 basepair reads (UCSF CAT).

**Table 3:**
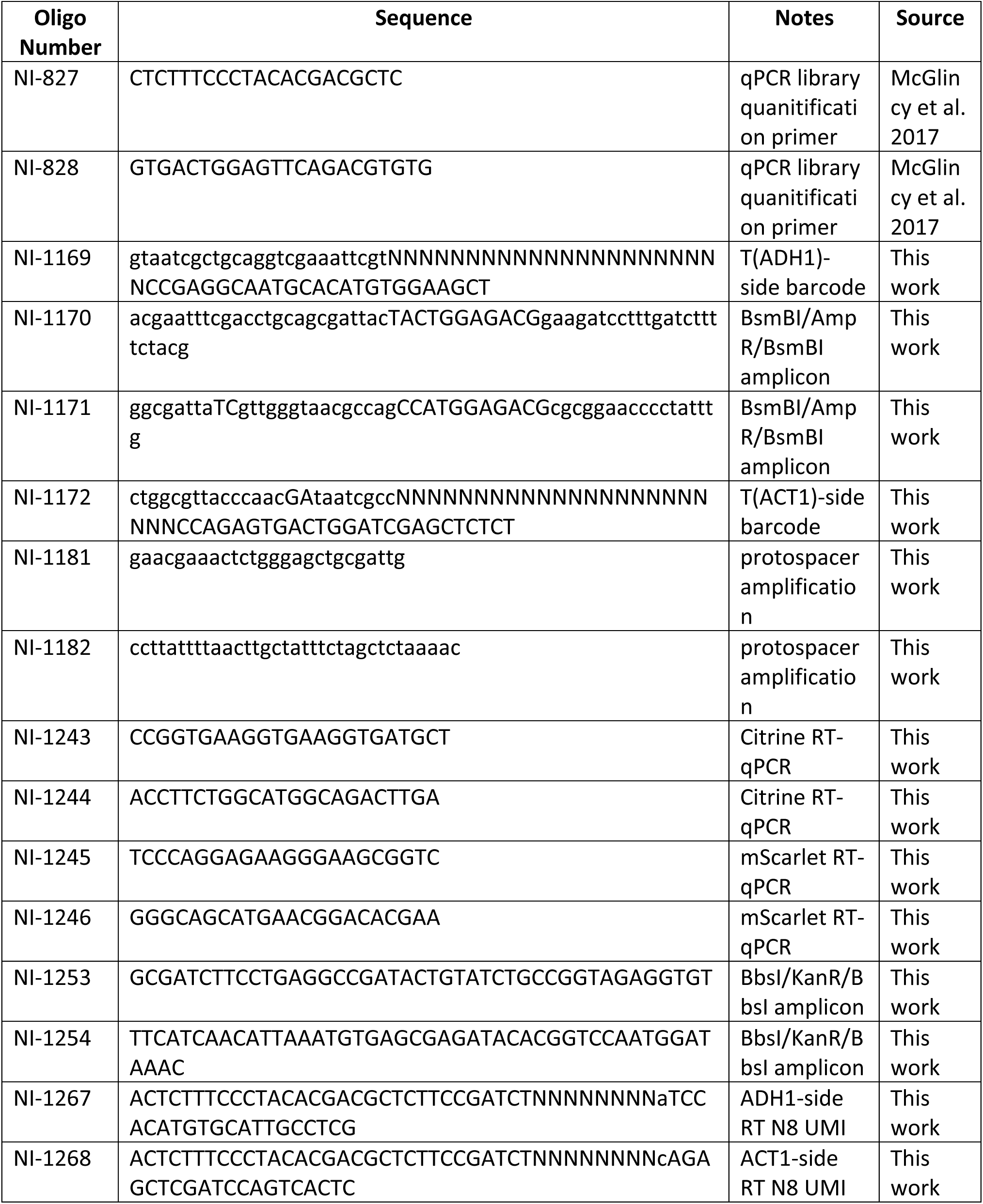
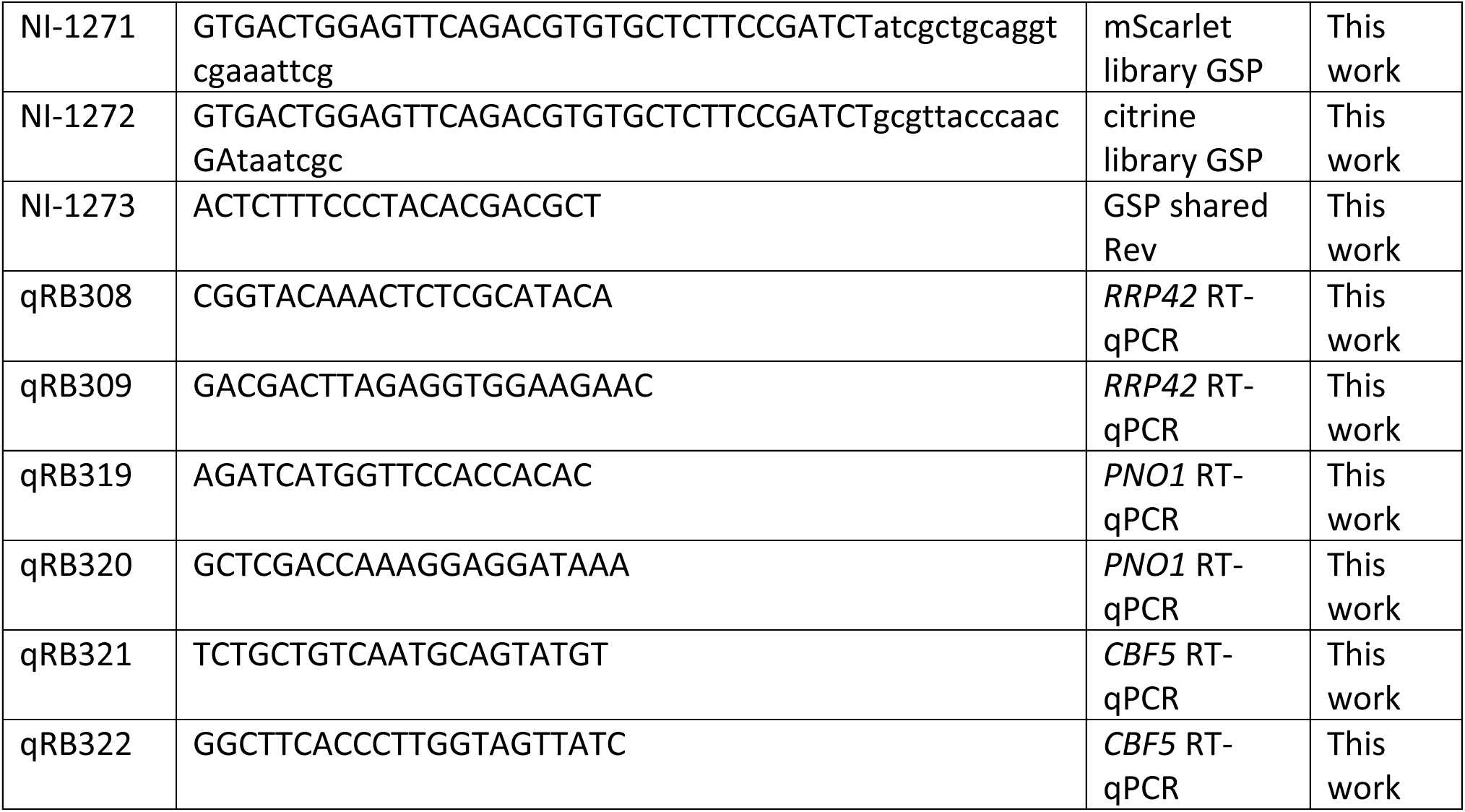
Primers used in this study.

Sequencing reads were trimmed using cutadapt to remove Illumina adapter sequences (Martin 2011). Samtools software was used to process and prepare the alignment and reads were counted using HTSeq-count (Li et al. 2009; Anders, Pyl, and Huber 2015). Differential expression analysis was performed with DESeq2 (Love, Huber, and Anders 2014). Filtered DESeq2 dataset (number of barcodes ≥2) is available in Table 5. Filtered DESeq2 dataset normalized to negative control guide and accompanying heatmap can be found in Table 6 and Supplementary File 1.

### Growth Assays

Cells were grown overnight in minimal media to saturation. Cells were diluted to OD_600_ of 0.2 and grown to mid-log phase (typically 5 mL culture at OD_600_ 0.5 - 1.0), and then further diluted to OD_600_ 0.01 in triplicate in a 96-well plate. To generate the SM dose response curve, SM was added to cells upon plate setup. For tetracycline dose response curve, tetracycline was added to cells upon plate setup. Absorbance measurements at OD_595_ were taken every 15 minutes, with shaking in between measurements, and temperature maintained at 30 °C (Tecan SPARK Multimode Microplate Reader). Growth rates were fit using the R package ‘growthcurver’ (Sprouffske and Wagner 2016).

### RT-qPCR

For guide validation, cells were grown to saturation overnight in minimal media. Cells were diluted to OD_600_ of 0.1 and grown at 30 °C to mid-log phase (OD_600_ 0.5-0.8). Cells were then further diluted to OD_600_ of 0.025 for 5 ml culture without tetracycline and 60 ml culture with 500 ng/ml tetracycline. After ∼17.5 hours, 5 ml culture without tetracycline was harvested as pre-induction sample and 5 ml culture with tetracycline was harvested as post-guide induction sample. Two aliquots of 25ml culture were split into two pre-warmed flasks and one flask was treated with low dose SM (0.56 µM) and one flask was treated with high dose SM (14 µM). Cells from each treatment dose were collected at .25 hours, 1 hour, and 4 hour timepoints.

RNA was extracted from cell pellets using phenol/chloroform method in (Ares 2012). Complementary DNA was synthesized using ProtoScript II reverse transcriptase (NEB #M0368L) and oligo d(T)_23_VN according to manufacturer’s protocol. Quantitative PCR was conducted using the Luna universal qPCR master mix (NEB #M3003L) and the primers listed in Table 3 (BioRad-CFX96 qPCR System instrument).

### Sucrose Gradients

Cells were grown to mid-log phase in a volume of at least 150 ml (OD_600_ ∼0.5 -1.0). Cells were harvested by vacuum filtration with 0.45 µM Whatman cellulose nitrate membrane filters (Fisher Scientific #09-744-75). Yeast cells were immediately scraped off with a pre-chilled metal spatula and plunged into a conical tube with holes poked into the cap and filled with liquid nitrogen. 2 ml of freshly prepared ice cold polysome buffer (20 mM Tris-HCl pH 7.4, 150 mM NaCl, 5 mM MgCl_2_, 1 mM DTT, 1% Triton X-100, .02 U/µl Turbo DNase) were dripped into the conical tube with cells, and frozen samples were stored at -80 °C.

Frozen yeast and polysome buffer were lysed by cryogrinding with the MM400 Mixer mill (Retsch #20.745.0001) for six cycles of 3 minutes at 15 Hz. The powder was moved from the grinding chamber using a pre-chilled metal spatula into a conical tube with holes poked into the cap, filled with liquid nitrogen, and stored at -80 °C overnight. Lysates were thawed slowly on an ice bucket (∼2 hours) and clarified by centrifuging the conical tube at 3,000 x g for 5 min at 4 °C. Supernatant was transferred to Eppendorf tubes and clarified by centrifugation at 14,000 rpm for 15 min at 4 °C. Supernatant was transferred to Eppendorf tubes. Lysate was quantified by isolating RNA from 100 µl using a Direct-zol RNA Purification Kit (Zymo #R2060) and treating with TurboDNase for 15 minutes. Clarified lysates were used for sucrose gradients.

Sucrose gradients were prepared using a Gradient Master (BioComp Instruments) using Seton Open-Top Polyclear Centrifuge Tubes (Seton Scientific #7030). Normalized lysates (220µl) were loaded onto a 10–50% sucrose gradient prepared in polysome buffer. Gradients were centrifuged at 35,000 rpm for 3 hours at 4 °C in a Beckman SW41 Ti rotor using an XL-70 ultracentrifuge (Beckman Coulter). The sucrose gradients were analyzed using the gradient master system and the absorbance was monitored using a UV monitor (BioRad EM-1 Econo UV monitor). Short density gradients followed the same protocol, except using 5–30% sucrose concentrations.

---

**Supplementary Figure 1:**
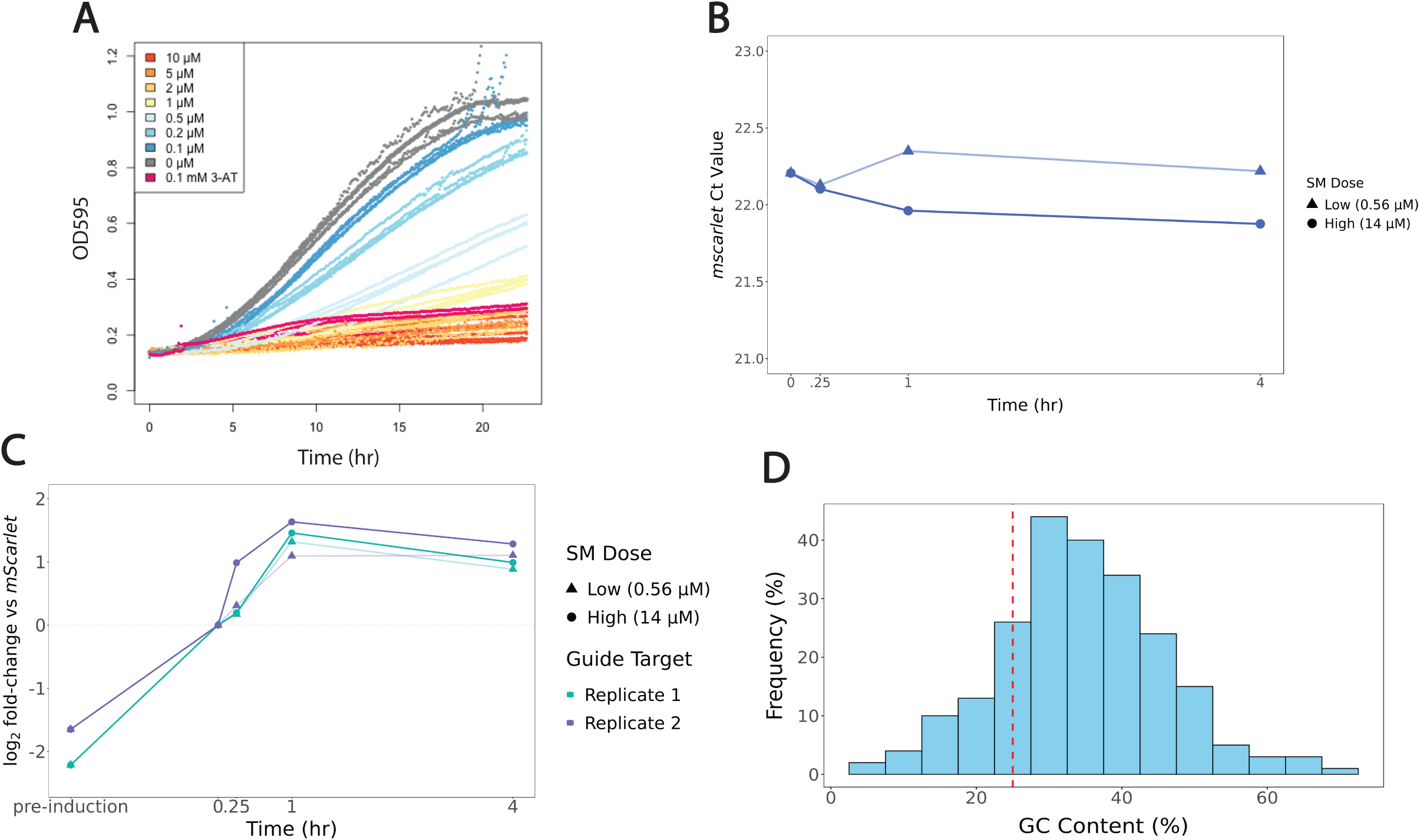
Additional pilot data (A) Raw data growth curves from Figure 4. 1A **(B)** RT-qPCR for *UBC6* driven mScarlet normalizer reporter shows minimal variability in Ct values for low SM dose (●) or high SM dose (▲) treated cells after 0, .25, 1, or 4 hours. **(C)** RT-qPCR of ISR reporter normalized to mScarlet for CiBER-seq libraries. Two biological replicates for low and high dose SM treatment. **(D)** Histogram of percentage of GC content in 224 CiBER-seq spacer sequences. Dashed line represents 25% cutoff mark; spacer sequences that fell below this threshold were not followed-up on due to increased likelihood of off-target effects (Konstantakos et al. 2022).

**Supplementary Figure 2:**
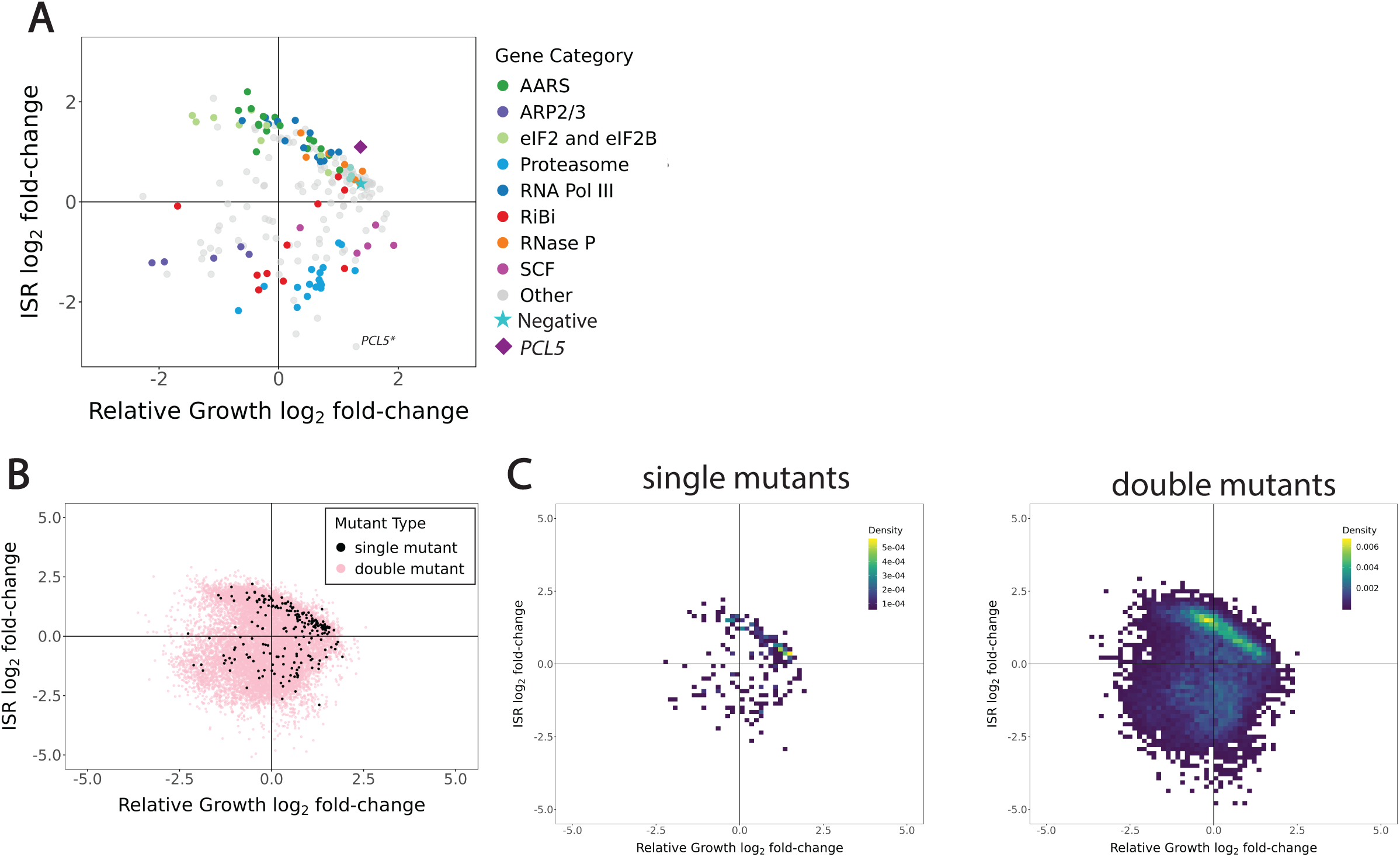
Categorized single mutant analysis and double mutant analysis. **(A)** Guides targeting functionally related genes show similar growth and ISR activation phenotypes after guide-induction. **(B)** Dual-guide phenotypes (pink) and single-guide phenotypes (black) from DESeq2 analysis **(C)** Single-guide (left) and double-guide density plots (right) show similar density patterns within the range of the single-mutant phenotypes. Many guides activate the ISR and cause cellular fitness defects in a proportionate way.

**Supplementary Figure 3:**
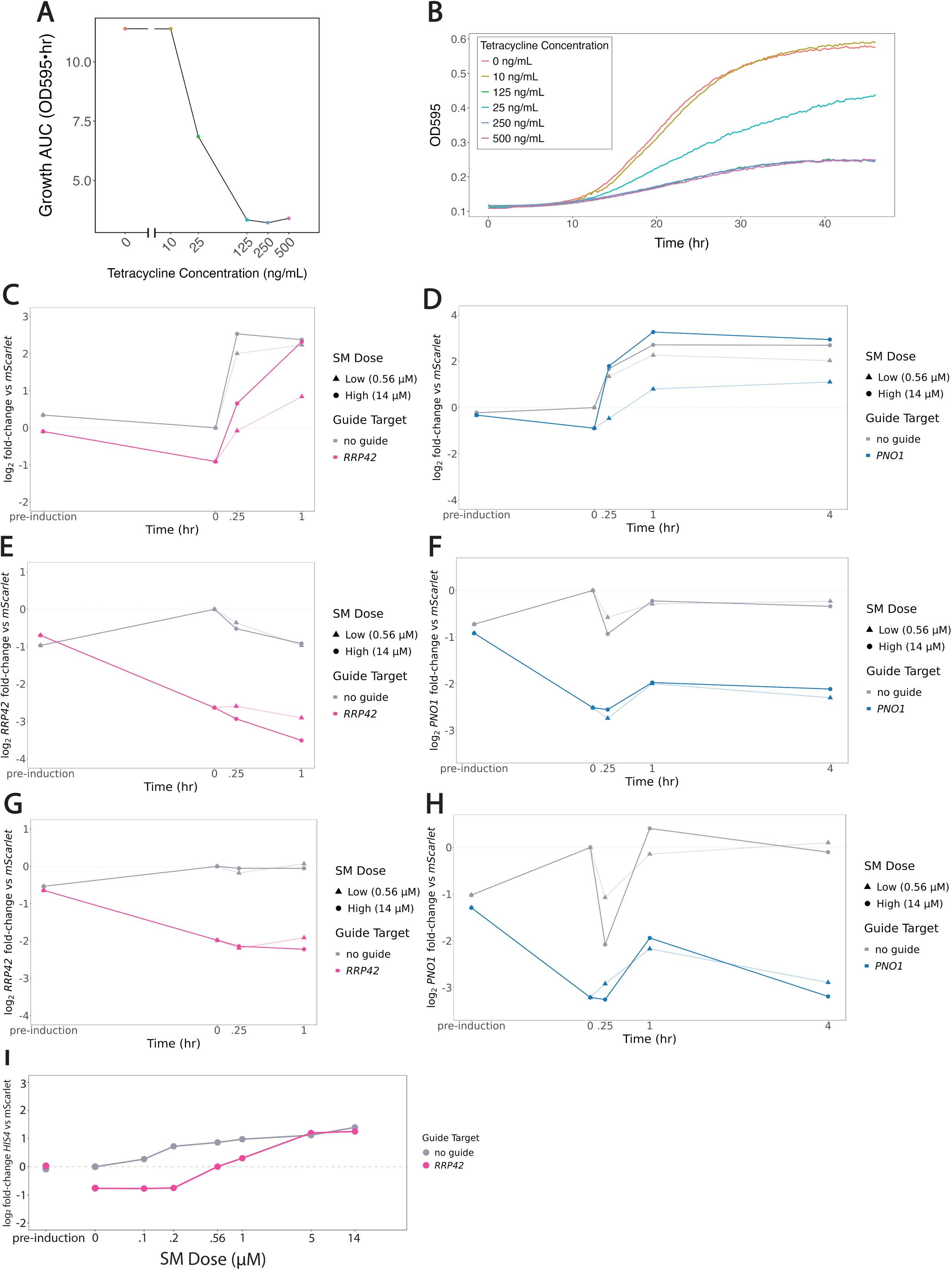
Replicates of *RRP42* and *PNO1* lead to SM treatment dose-dependent response. **(A)** Growth AUC (Area Under Logistic Curve) reflects relative growth rate for fitted growth curves for tetracycline guide-induction dose response. Growth rate slows with increased knockdown efficiency. **(B)** Raw growth curves from (A). **(C)** RT-qPCR of ISR reporter for wildtype and *RRP42* knockdown cells, replicate 2 of Figure 3C. **(D)** RT-qPCR of ISR reporter for wildtype and *PNO1* knockdown cells, replicate 2 of Figure 3D. **(E)** RT-qPCR *RRP42* knockdown-efficiency of data from Figure 3C **(F)** RT-qPCR *PNO1* knockdown-efficiency of Figure 3D. **(G)** RT-qPCR *RRP42* knockdown-efficiency of data from Supp Figure 3C. **(H)** RT-qPCR *PNO1* knockdown-efficiency of data from Supp Figure 3D. **(I)** RT-qPCR for endogenous *HIS4* target for SM dose response from Figure 3E. HIS4 expression in *RRP42* knockdown cells shows similar decrease to ISR reporter.

**Supplementary Figure 4:**
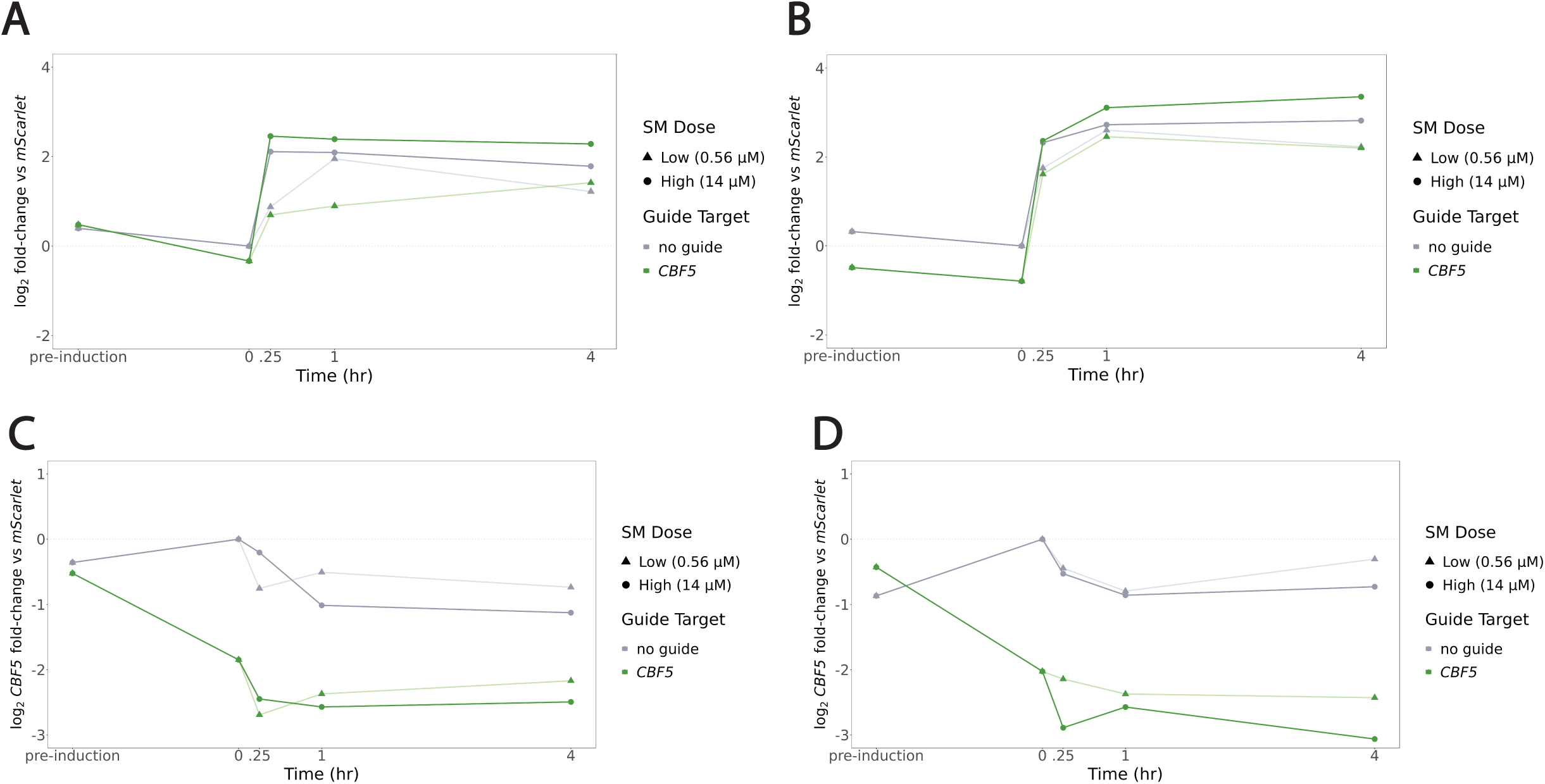
Knockdown of RiBi gene *CBF5* SM dose-dependent effect. **(A)** RT-qPCR of ISR reporter for wildtype and *PNO1* knockdown cells, replicate 1. ISR reporter normalized to mScarlet for pre-guide induction, post-guide induction and 0, .25, 1 and 4 hour low or high SM treated cells. Low SM dose reduces ISR activation and high SM dose strengthens ISR activation. **(B)** RT-qPCR of ISR reporter for wildtype and *CBF5* knockdown cells, replicate 2 of Supp Figure 4B. **(C)** RT-qPCR *CBF5* knockdown-efficiency of data from Supp Figure 4A. **(D)** RT-qPCR *CBF5* knockdown-efficiency of data from Supp Figure 4B.

**Supplementary Figure 5:**
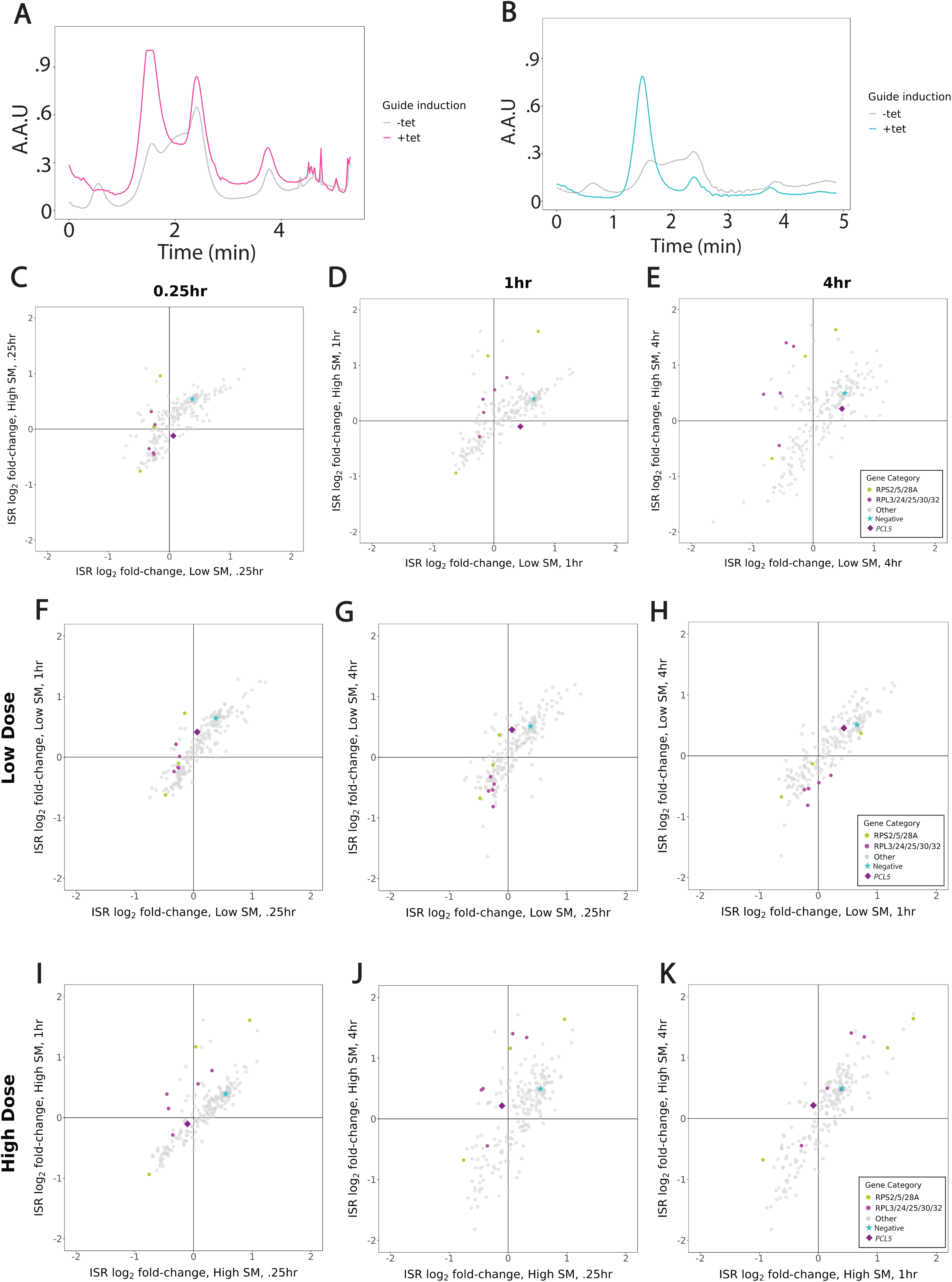
Targeting ribosomal genes effects ISR dynamics. **(A-B)** Short 5-30% sucrose gradients for **(A)** *RRP42* knockdown and **(B)** *PNO1* knockdown lysates. **(C-E)** Scatterplots highlighting guides targeting small and large ribosome subunits for low versus high SM treatment dose for **(C)** .25 hour **(D)** 1 hour **(E)** 4 hours. (F-H) Scatterplots highlighting guides targeting small and large ribosome subunits for low dose SM treatment **(F)** .25hr versus 1hr timepoints **(G)** .25hr versus 4hr timepoints **(H)** 1hr versus 4 hour timepoints. **(I-K)** Scatterplots highlighting guides targeting small and large ribosome subunits for high dose SM treatment **(I)** .25hr versus 1hr timepoints **(J)** .25hr versus 4hr timepoints **(K)** 1hr versus 4 hour timepoints.

## References

1. Abdulkarim, Baroj, Marc Nicolino, Mariana Igoillo-Esteve, Mathilde Daures, Sophie Romero, Anne Philippi, Valérie Senée, et al. 2015. “A Missense Mutation in PPP1R15B Causes a Syndrome Including Diabetes, Short Stature, and Microcephaly.” Diabetes 64 (11): 3951–62. 10.2337/db15-0477.

2. Albert, Benjamin, Isabelle C Kos-Braun, Anthony K Henras, Christophe Dez, Maria Paula Rueda, Xu Zhang, Olivier Gadal, Martin Kos, and David Shore. 2019. “A Ribosome Assembly Stress Response Regulates Transcription to Maintain Proteome Homeostasis.” Edited by Naama Barkai, Alan G Hinnebusch, John Woolford, and Marlene Oeffinger. eLife 8 (May):e45002. 10.7554/eLife.45002.

3. Alford, Brian D., Eduardo Tassoni-Tsuchida, Danish Khan, Jeremy J. Work, Gregory Valiant, and Onn Brandman. 2021. “ReporterSeq Reveals Genome-Wide Dynamic Modulators of the Heat Shock Response across Diverse Stressors.” eLife 10 (July):e57376. 10.7554/eLife.57376.

4. Anders, Simon, Paul Theodor Pyl, and Wolfgang Huber. 2015. “HTSeq—a Python Framework to Work with High-Throughput Sequencing Data.” Bioinformatics 31 (2): 166–69. 10.1093/bioinformatics/btu638.

5. Ares, Manuel. 2012. “Isolation of Total RNA from Yeast Cell Cultures.” Cold Spring Harbor Protocols 2012 (10): 1082–86. 10.1101/pdb.prot071456.

6. Cheng, Ze, Christopher Frederick Mugler, Abdurrahman Keskin, Stefanie Hodapp, Leon Yen-Lee Chan, Karsten Weis, Philipp Mertins, Aviv Regev, Marko Jovanovic, and Gloria Ann Brar. 2019. “Small and Large Ribosomal Subunit Deficiencies Lead to Distinct Gene Expression Signatures That Reflect Cellular Growth Rate.” Molecular Cell 73 (1): 36–47.e10. 10.1016/j.molcel.2018.10.032.

7. Costa-Mattioli, Mauro, and Peter Walter. 2020. “The Integrated Stress Response: From Mechanism to Disease.” Science 368 (6489): eaat5314. 10.1126/science.aat5314.

8. Das, Indrajit, Agnieszka Krzyzosiak, Kim Schneider, Lawrence Wrabetz, Maurizio D’Antonio, Nicholas Barry, Anna Sigurdardottir, and Anne Bertolotti. 2015. “Preventing Proteostasis Diseases by Selective Inhibition of a Phosphatase Regulatory Subunit.” Science (New York, N.Y.) 348 (6231): 239–42. 10.1126/science.aaa4484.

9. Delépine, M., M. Nicolino, T. Barrett, M. Golamaully, G. M. Lathrop, and C. Julier. 2000. “EIF2AK3, Encoding Translation Initiation Factor 2-Alpha Kinase 3, Is Mutated in Patients with Wolcott-Rallison Syndrome.” Nature Genetics 25 (4): 406–9. 10.1038/78085.

10. Diamond, Paige D., Nicholas J. McGlincy, and Nicholas T. Ingolia. 2024. “Depletion of Cap-Binding Protein eIF4E Dysregulates Amino Acid Metabolic Gene Expression.” Molecular Cell 84 (11): 2119–2134.e5. 10.1016/j.molcel.2024.05.008.

11. Durrant, Matthew G., Alison Fanton, Josh Tycko, Michaela Hinks, Sita S. Chandrasekaran, Nicholas T. Perry, Julia Schaepe, et al. 2023. “Systematic Discovery of Recombinases for Efficient Integration of Large DNA Sequences into the Human Genome.” Nature Biotechnology 41 (4): 488–99. 10.1038/s41587-022-01494-w.

12. Gibson, Daniel G., Lei Young, Ray-Yuan Chuang, J. Craig Venter, Clyde A. Hutchison, and Hamilton O. Smith. 2009. “Enzymatic Assembly of DNA Molecules up to Several Hundred Kilobases.” Nature Methods 6 (5): 343–45. 10.1038/nmeth.1318.

13. Gietz, R. Daniel, and Robert H. Schiestl. 2007. “High-Efficiency Yeast Transformation Using the LiAc/SS Carrier DNA/PEG Method.” Nature Protocols 2 (1): 31–34. 10.1038/nprot.2007.13.

14. Gilbert, Luke A., Matthew H. Larson, Leonardo Morsut, Zairan Liu, Gloria A. Brar, Sandra E. Torres, Noam Stern-Ginossar, et al. 2013. “CRISPR-Mediated Modular RNA-Guided Regulation of Transcription in Eukaryotes.” Cell 154 (2): 442–51. 10.1016/j.cell.2013.06.044.

15. Halliday, M., H. Radford, Y. Sekine, J. Moreno, N. Verity, J. le Quesne, C. A. Ortori, et al. 2015. “Partial Restoration of Protein Synthesis Rates by the Small Molecule ISRIB Prevents Neurodegeneration without Pancreatic Toxicity.” Cell Death & Disease 6 (3): e1672. 10.1038/cddis.2015.49.

16. Harding, H. P., Y. Zhang, A. Bertolotti, H. Zeng, and D. Ron. 2000. “Perk Is Essential for Translational Regulation and Cell Survival during the Unfolded Protein Response.” Molecular Cell 5 (5): 897–904. 10.1016/s1097-2765(00)80330-5.

17. Harding, Heather P., Yuhong Zhang, Huiquing Zeng, Isabel Novoa, Phoebe D. Lu, Marcella Calfon, Navid Sadri, et al. 2003. “An Integrated Stress Response Regulates Amino Acid Metabolism and Resistance to Oxidative Stress.” Molecular Cell 11 (3): 619–33. 10.1016/s1097-2765(03)00105-9.

18. Hinnebusch, Alan G. 2005. “Translational Regulation of GCN4 and the General Amino Acid Control of Yeast.” Annual Review of Microbiology 59 (1): 407–50. 10.1146/annurev.micro.59.031805.133833.

19. Jia, M. H., R. A. Larossa, J. M. Lee, A. Rafalski, E. Derose, G. Gonye, and Z. Xue. 2000. “Global Expression Profiling of Yeast Treated with an Inhibitor of Amino Acid Biosynthesis, Sulfometuron Methyl.” Physiological Genomics 3 (2): 83–92. 10.1152/physiolgenomics.2000.3.2.83.

20. Kernohan, Kristin D., Martine Tétreault, Urszula Liwak-Muir, Michael T. Geraghty, Wen Qin, Sunita Venkateswaran, Jorge Davila, et al. 2015. “Homozygous Mutation in the Eukaryotic Translation Initiation Factor 2alpha Phosphatase Gene, PPP1R15B, Is Associated with Severe Microcephaly, Short Stature and Intellectual Disability.” Human Molecular Genetics 24 (22): 6293–6300. 10.1093/hmg/ddv337.

21. Kilchert, Cornelia, Sina Wittmann, and Lidia Vasiljeva. 2016. “The Regulation and Functions of the Nuclear RNA Exosome Complex.” Nature Reviews. Molecular Cell Biology 17 (4): 227–39. 10.1038/nrm.2015.15.

22. Kim, Jinyoung, Ryan Y. Muller, Eliana R. Bondra, and Nicholas T. Ingolia. 2024. “CRISPRi with Barcoded Expression Reporters Dissects Regulatory Networks in Human Cells.” bioRxiv: The Preprint Server for Biology, September, 2024.09.06.611573. 10.1101/2024.09.06.611573.

23. Klein, Philipp, Stefan M. Kallenberger, Hanna Roth, Karsten Roth, Thi Bach Nga Ly-Hartig, Vera Magg, Janez Aleš, et al. 2022. “Temporal Control of the Integrated Stress Response by a Stochastic Molecular Switch.” Science Advances 8 (12): eabk2022. 10.1126/sciadv.abk2022.

24. Lafontaine, D. L., C. Bousquet-Antonelli, Y. Henry, M. Caizergues-Ferrer, and D. Tollervey. 1998. “The Box H + ACA snoRNAs Carry Cbf5p, the Putative rRNA Pseudouridine Synthase.” Genes & Development 12 (4): 527–37. 10.1101/gad.12.4.527.

25. Lee, Kihoon, Yu Zhang, and Sang Eun Lee. 2008. “Saccharomyces Cerevisiae ATM Orthologue Suppresses Break-Induced Chromosome Translocations.” Nature 454 (7203): 543–46. 10.1038/nature07054.

26. Lee, Michael E., William C. DeLoache, Bernardo Cervantes, and John E. Dueber. 2015. “A Highly Characterized Yeast Toolkit for Modular, Multipart Assembly.” ACS Synthetic Biology 4 (9): 975–86. 10.1021/sb500366v.

27. Levy, Sasha F., Jamie R. Blundell, Sandeep Venkataram, Dmitri A. Petrov, Daniel S. Fisher, and Gavin Sherlock. 2015. “Quantitative Evolutionary Dynamics Using High-Resolution Lineage Tracking.” Nature 519 (7542): 181–86. 10.1038/nature14279.

28. Li, Heng, Bob Handsaker, Alec Wysoker, Tim Fennell, Jue Ruan, Nils Homer, Gabor Marth, Goncalo Abecasis, and Richard Durbin. 2009. “The Sequence Alignment/Map Format and SAMtools.” Bioinformatics 25 (16): 2078–79. 10.1093/bioinformatics/btp352.

29. Lobel, Joseph H., and Nicholas T. Ingolia. 2024. “Precise Measurement of Molecular Phenotypes with Barcode-Based CRISPRi Systems.” bioRxiv: The Preprint Server for Biology, June, 2024.06.21.600132. 10.1101/2024.06.21.600132.

30. Love, Michael I., Wolfgang Huber, and Simon Anders. 2014. “Moderated Estimation of Fold Change and Dispersion for RNA-Seq Data with DESeq2.” Genome Biology 15 (12): 550. 10.1186/s13059-014-0550-8.

31. Ma, Yanjun, and Linda M. Hendershot. 2003. “Delineation of a Negative Feedback Regulatory Loop That Controls Protein Translation during Endoplasmic Reticulum Stress.” The Journal of Biological Chemistry 278 (37): 34864–73. 10.1074/jbc.M301107200.

32. Martin, Marcel. 2011. “Cutadapt Removes Adapter Sequences from High-Throughput Sequencing Reads.” EMBnet.Journal 17 (1): 10–12. 10.14806/ej.17.1.200.

33. Matreyek, Kenneth A., Jason J. Stephany, Melissa A. Chiasson, Nicholas Hasle, and Douglas M. Fowler. 2020. “An Improved Platform for Functional Assessment of Large Protein Libraries in Mammalian Cells.” Nucleic Acids Research 48 (1): e1. 10.1093/nar/gkz910.

34. Matreyek, Kenneth A., Jason J. Stephany, and Douglas M. Fowler. 2017. “A Platform for Functional Assessment of Large Variant Libraries in Mammalian Cells.” Nucleic Acids Research 45 (11): e102. 10.1093/nar/gkx183.

35. Mülleder, Michael, Floriana Capuano, Pınar Pir, Stefan Christen, Uwe Sauer, Stephen G Oliver, and Markus Ralser. 2012. “A Prototrophic Deletion Mutant Collection for Yeast Metabolomics and Systems Biology.” Nature Biotechnology 30 (12): 1176–78. 10.1038/nbt.2442.

36. Muller, Ryan, Zuriah A. Meacham, Lucas Ferguson, and Nicholas T. Ingolia. 2020. “CiBER-Seq Dissects Genetic Networks by Quantitative CRISPRi Profiling of Expression Phenotypes.” Science 370 (6522): eabb9662. 10.1126/science.abb9662.

37. Novoa, Isabel, Yuhong Zhang, Huiqing Zeng, Rivka Jungreis, Heather P. Harding, and David Ron. 2003. “Stress-induced Gene Expression Requires Programmed Recovery from Translational Repression.” The EMBO Journal 22 (5): 1180–87. 10.1093/emboj/cdg112.

38. Okuda, Ellen K., Fernando A. Gonzales-Zubiate, Olivier Gadal, and Carla C. Oliveira. 2020. “Nucleolar Localization of the Yeast RNA Exosome Subunit Rrp44 Hints at Early Pre-rRNA Processing as Its Main Function.” The Journal of Biological Chemistry 295 (32): 11195–213. 10.1074/jbc.RA120.013589.

39. Scheuner, D., B. Song, E. McEwen, C. Liu, R. Laybutt, P. Gillespie, T. Saunders, S. Bonner-Weir, and R. J. Kaufman. 2001. “Translational Control Is Required for the Unfolded Protein Response and in Vivo Glucose Homeostasis.” Molecular Cell 7 (6): 1165–76. 10.1016/s1097-2765(01)00265-9.

40. Senapin, Saengchan, G. Desmond Clark-Walker, Xin Jie Chen, Bertrand Séraphin, and Marie-Claire Daugeron. 2003. “RRP20, a Component of the 90S Preribosome, Is Required for Pre-18S rRNA Processing in Saccharomyces Cerevisiae.” Nucleic Acids Research 31 (10): 2524–33. 10.1093/nar/gkg366.

41. Shemer, Revital, Ariella Meimoun, Tsvi Holtzman, and Daniel Kornitzer. 2002. “Regulation of the Transcription Factor Gcn4 by Pho85 Cyclin PCL5.” Molecular and Cellular Biology 22 (15): 5395–5404. 10.1128/MCB.22.15.5395-5404.2002.

42. Sidrauski, Carmela, Diego Acosta-Alvear, Arkady Khoutorsky, Punitha Vedantham, Brian R Hearn, Han Li, Karine Gamache, et al. 2013. “Pharmacological Brake-Release of mRNA Translation Enhances Cognitive Memory.” Edited by David Ron. eLife 2 (May):e00498. 10.7554/eLife.00498.

43. Smith, Justin D., Sundari Suresh, Ulrich Schlecht, Manhong Wu, Omar Wagih, Gary Peltz, Ronald W. Davis, Lars M. Steinmetz, Leopold Parts, and Robert P. St.Onge. 2016. “Quantitative CRISPR Interference Screens in Yeast Identify Chemical-Genetic Interactions and New Rules for Guide RNA Design.” Genome Biology 17 (1): 45. 10.1186/s13059-016-0900-9.

44. Sprouffske, Kathleen, and Andreas Wagner. 2016. “Growthcurver: An R Package for Obtaining Interpretable Metrics from Microbial Growth Curves.” BMC Bioinformatics 17 (April):172. 10.1186/s12859-016-1016-7.

45. Stovicek, Vratislav, Gheorghe M. Borja, Jochen Forster, and Irina Borodina. 2015. “EasyClone 2.0: Expanded Toolkit of Integrative Vectors for Stable Gene Expression in Industrial Saccharomyces Cerevisiae Strains.” Journal of Industrial Microbiology & Biotechnology 42 (11): 1519–31. 10.1007/s10295-015-1684-8.

46. Tye, Blake W, Nicoletta Commins, Lillia V Ryazanova, Martin Wühr, Michael Springer, David Pincus, and L Stirling Churchman. 2019. “Proteotoxicity from Aberrant Ribosome Biogenesis Compromises Cell Fitness.” Edited by Alan G Hinnebusch, Naama Barkai, and David Tollervey. eLife 8 (March):e43002. 10.7554/eLife.43002.

47. Wu, Colin Chih-Chien, Amy Peterson, Boris Zinshteyn, Sergi Regot, and Rachel Green. 2020. “Ribosome Collisions Trigger General Stress Responses to Regulate Cell Fate.” Cell 182 (2): 404–416.e14. 10.1016/j.cell.2020.06.006.

48. Xu, Zhengyao, and William R. A. Brown. 2016. “Comparison and Optimization of Ten Phage Encoded Serine Integrases for Genome Engineering in Saccharomyces Cerevisiae.” BMC Biotechnology 16 (1): 13. 10.1186/s12896-016-0241-5.

