## Supplementary File 1 for "Mapping the Genetic Architecture of the Adaptive Integrated Stress Response in *S. cerevisiae*"

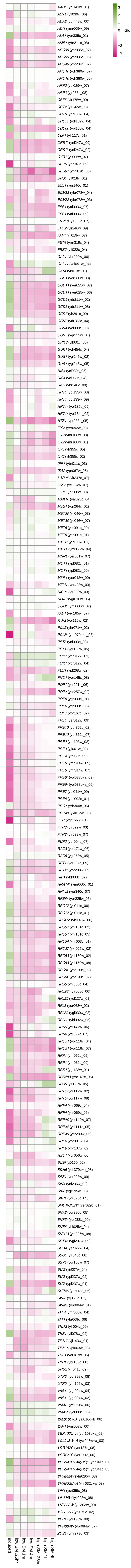


**Supplementary File 1:** Heatmap of all single mutants shows ISR log2 fold-change normalized to negative guide reference. Guides marked with a * reflect updated guide targets at divergent promoters that were misassigned in the original table.
